# Exploiting pair correlation function to describe biological tissue structure

**DOI:** 10.64898/2025.12.19.695425

**Authors:** Fatoumata Mangane, Ruben Casanova, Kyra JE Borgman, Leanne De Koning, Martina Haberecker, Chantal Pauli, Susanne Dettwiler, Sergio Roman-Roman, Bernd Bodenmiller, Manuel Rodrigues, Pierre Bost

## Abstract

Multiplexed imaging technologies now enable the simultaneous profiling of hundreds to thousands of molecular targets in intact tissues, providing unprecedented insight into cellular heterogeneity and spatial organization. While data generation has rapidly matured, the quantitative analysis of spatial structure remains challenging and poorly standardized, particularly across biological length scales. Existing approaches, such as distance-based metrics, neighborhood analyses and graph neural networks, either capture only local interactions or sacrifice interpretability for predictive power. Here we introduce **PCF-SiM** (Pair Correlation Function Sigmoid Modeling), a scalable and interpretable framework that leverages parametric modeling of the pair correlation function to quantify spatial organization in multiplexed imaging data. PCF-SiM compresses complex spatial patterns into a small set of biologically meaningful parameters, enabling robust comparisons across cell types, samples and conditions. Applying PCF-SiM to diverse public spatial transcriptomics datasets, we demonstrate its ability to detect condition-dependent tissue remodeling in a mouse colitis model. We further extend the framework with a co-scaling strategy that identifies cell types participating in shared spatial structures. Using newly generated clinical datasets from Hashimoto’s thyroiditis and uveal melanoma liver metastases, PCF-SiM reveals hierarchical organization of autoimmune infiltrates and coordinated spatial interactions between lymphatic endothelial cells and tumor-infiltrating lymphocytes. Finally, we show that reliable inference of tissue-scale architecture requires whole-slide imaging, exposing intrinsic limitations of tumor microarray–based spatial analyses. Together, PCF-SiM provides a principled and interpretable approach for spatial analysis of multiplexed imaging data.

## Introduction

Multiplexed imaging (MI) technologies comprise a class of advanced microscopy-based methods that enable the simultaneous measurement of hundreds to thousands of gene or protein targets directly in intact tissues. These approaches typically rely on iterative cycles of staining, imaging, and signal removal to accumulate molecular information with high spatial precision [Moffit et al., 2022]. In contrast to dissociative single-cell RNA sequencing (scRNA-seq), MI preserves the native spatial organization of cells, allowing researchers to study cellular heterogeneity together with cell–cell interactions and tissue architecture. Over the last decade, continuous technological improvements have substantially expanded the capabilities of MI platforms [Moses and Pachter, 2022]. Current systems can now profile thousands of transcripts and hundreds of proteins in parallel with subcellular resolution and high sensitivity. Importantly, the recent commercialization of RNA-based MI platforms such as 10x Genomics Xenium [Janesick et al., 2022] and NanoString CosMx [He et al., 2022] has made these technologies broadly accessible to non-specialist laboratories, accelerating their adoption across biomedical research. These platforms have already enabled high-resolution spatial atlases of both healthy tissues, such as mouse brain [Yao et al., 2023], human brain [Qian et al., 2025], and human liver [Watson et al., 2025], and pathological contexts, including human breast tumors [Jackson et al., 2020], fibrotic liver [Watson et al., 2025], and injured mouse heart tissue [Shing-Fung Chan et al., 2025].

While the generation of high-quality spatial data is now within reach of many laboratories, the analysis of such data remains challenging and insufficiently standardized, particularly when it comes to quantifying spatial organization. Currently, MI-derived spatial data are typically analyzed using three main approaches: (i) distance-based analyses, (ii) neighborhood-based analyses, and (iii) graph neural network (GNN) methods. Distance-based approaches examine metrics such as the minimum or average distance between cells of the same or different types. In contrast, neighborhood-based analyses characterize the cellular composition surrounding each cell, often identifying recurrent or “archetypal” spatial neighborhoods [Schürch et al., 2020]. GNN-based methods construct a graph where nodes represent cells and edges denote spatial proximity; these graphs are then used as input to neural networks to predict sample-level phenotypes [Ali et al., 2025]. Although all three approaches have been widely used, they each present limitations. Distance– and neighborhood-based methods primarily capture local spatial relationships and therefore provide limited insight into larger-scale spatial structure. Conversely, GNN approaches typically require large sample sizes for reliable training and are often perceived as “black boxes,” making their outputs more difficult to interpret. These limitations highlight the need for methods that are both interpretable and capable of capturing spatial structure across multiple biological scales.

A promising source of inspiration for developing new multiplexed imaging (MI) data analysis tools is the field of point pattern analysis [Illian et al., 2008; Bost et al., 2025], which has a long history of applications in spatial ecology and forestry. In particular, functional summary statistics such as Ripley’s K-function, Besag’s L-function, and the pair correlation function (pcf) are of high interest, as they have been widely used to characterize spatial organization across many scientific contexts. Briefly, the K-function describes, for a given radius r, the expected number of points within a circle of radius r centered on an arbitrary point. The L-function is a variance-stabilizing square-root transform of the K-function, while the pcf is defined as the derivative of the K-function with respect to r, normalized by 2πr, and can be interpreted as the expected number of additional points in an infinitesimally thin ring at distance r. The pcf has several advantages over the K– and L-functions. First, its values are directly interpretable: for a given distance r, pcf values greater than 1 indicate spatial aggregation, values equal to 1 correspond to spatial randomness, and values below 1 indicate repulsion [Illian et al., 2008]. Second, because it is non-cumulative, each value of the pcf reflects spatial structure specifically at that distance, without being influenced by patterns at smaller scales. Despite these advantages, the pcf is still rarely used in practice, largely because it produces a full function rather than a single scalar value for each cell type and sample, which makes downstream analyses, such as clustering samples or comparing cell types and sample groups, considerably more challenging.

An interesting strategy to facilitate the analysis of empirical pcfs is to summarize them into a small set of scalar descriptors by fitting a parametric function with a limited number of parameters (≤ 4). This approach is conceptually similar to that used for variograms, another family of spatial statistics commonly applied to continuous random fields [Matheron, 1963; Cressie, 1985]. However, because empirical pcfs and variograms often exhibit markedly different shapes, the functional models traditionally used to fit variograms are generally unsuitable for pcfs. To our knowledge, the parametric fitting of empirical pcfs has so far been explored only in astronomy, where the spatial distribution of galaxies is well described by a power-law model with an exponent between 1.5 and 1.9 [Snethlage et al., 2002]. Nevertheless, there is no guarantee that such a model is appropriate for other types of spatial data, particularly for biological tissues, whose organization arises from fundamentally different processes. Fitting empirical pcfs with a power-law or alternative functional families, therefore represents a promising strategy to develop robust and interpretable analysis tools for MI-derived spatial data.

To this end, we developed the Pair Correlation Function Sigmoid Modeling (PCF-SiM) framework, which transforms the spatial coordinates of cells within a tissue into a compact set of interpretable numerical parameters. After validating this approach across diverse publicly available spatial transcriptomics datasets, we first demonstrated that PCF-SiM enables robust differential structure analysis through the reanalysis of a colitis mouse model, revealing condition-dependent changes in tissue architecture. We then expanded the framework with a co-scaling strategy that identifies cell types participating in shared biological structures by detecting significant parameter covariation across samples. Leveraging this extended toolbox, we generated and analyzed two new clinical spatial datasets, Hashimoto’s thyroiditis and uveal melanoma liver metastasis, where PCF-SiM uncovered a hierarchical organization of the autoimmune infiltrate and a spatial interplay between lymphatic endothelial cells and tumor-infiltrating lymphocytes, respectively. Finally, we show that accurate structural inference requires whole-slide imaging rather than tumor microarrays, highlighting fundamental limitations of TMA-based spatial analyses.

## Results

### Fitting a sigmoid function is an efficient and easily interpretable pcf summarization approach

In order to assess the feasibility of studying cellular spatial structures through the fitting of pair correlation functions, we decided to process a set of publicly available spatial omic datasets generated using different tissues and technological platforms before computing the pcf of each individual identified cell type and then identify the function that fits best (Figure 1A). Regarding the datasets, we used a human lymph node dataset generated by Imaging Mass Cytometry (IMC) [Bost et al. 2023], another human lymph node dataset generated using the Xenium platform [Janesick et al. 2023], a human tonsil dataset generated by the MERFISH platform [Xia et al. 2019] and a CosMX [He et al. 2022] dataset derived from a human prefrontal cortex sample. While cell annotations were already available for the IMC datasets, this was not the case for the three other MI datasets. We thus used our recently published TranspaceR pipeline [Mangane et al. 2025] to analyze each dataset, resulting in cellular annotations in agreement with the cell types expected to be found in each sample (Supplementary Figure 1A, B and C).

**Figure 1.**
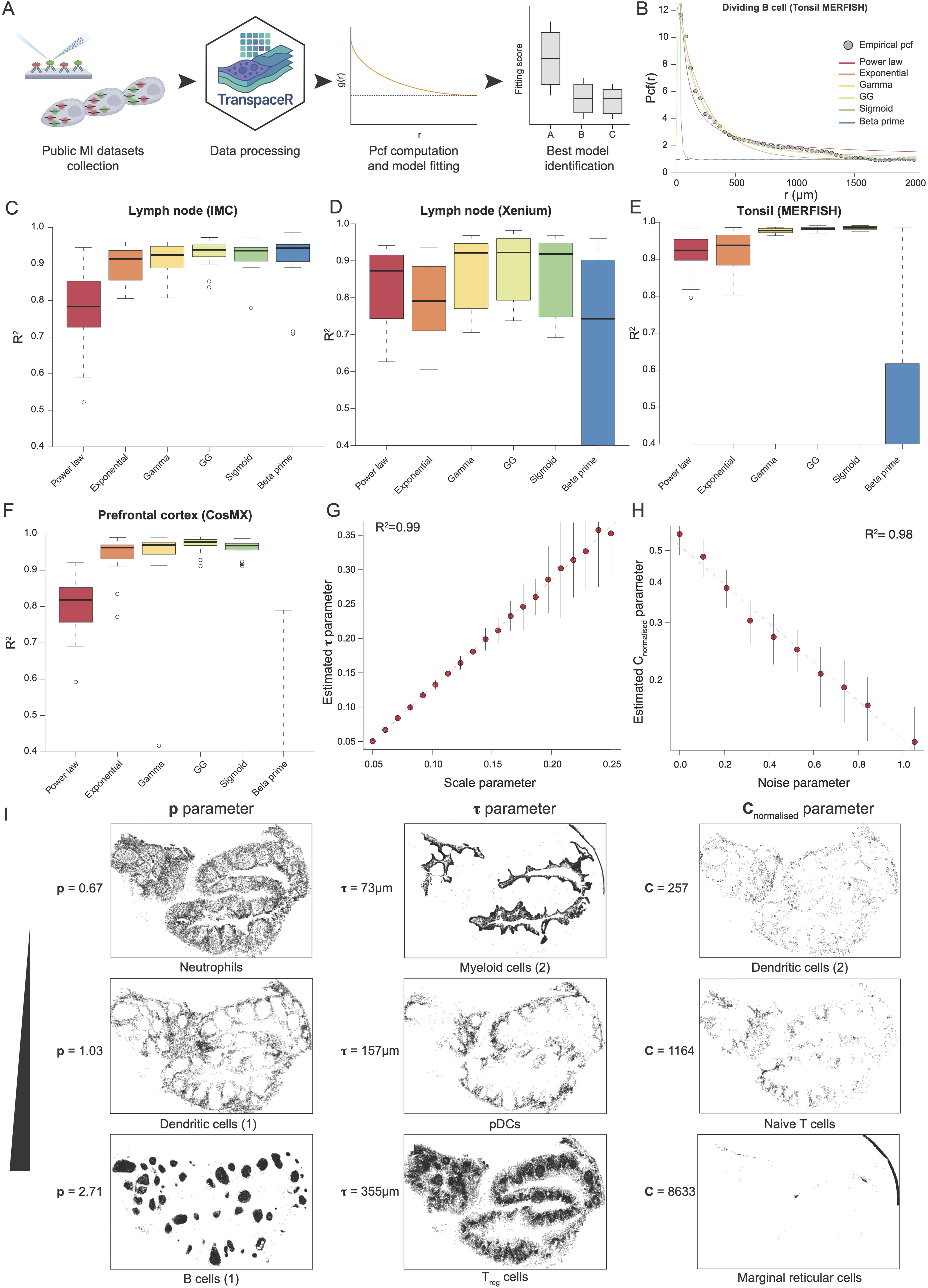
Identification of the sigmoid model for optimal pcf fitting. **(A)** Analytical workflow. **(B)** Example of pcf fit. Grey dots correspond to the empirical pcf. Each colored curve corresponds to the fit of a different model. The dashed line corresponds to the line of equation y=1. **(C)** Distribution of R^2^ values for the different model fits to the IMC Lymph node dataset. The thick line corresponds to the median, and the bottom and upper limits of the box correspond to the first and third quartiles, respectively. The lower and upper whiskers correspond to the lowest and highest values, respectively, within the range of the first and third quartiles ±1.5 times the interquartile range (IQR). All boxplots shown in this manuscript are using the same graphical code. The boxplot is based on N = 17 cell types. **(D)** Distribution of R^2^ values for the different model fit to the Xenium Lymph node dataset. The boxplot is based on N = 18 cell types. **(E)** Distribution of R^2^ values for the different model fit to the MERIFSH Tonsil dataset. The boxplot is based on N = 24 cell types. **(F)** Distribution of R^2^ values for the different model fit to the CosMX Prefrontal cortex dataset. The boxplot is based on N = 14 cell types. (**G**) Comparison of the inferred **τ** parameter with the scale parameter of the simulated Thomas point pattern. Each dot is the average result of 50 simulations. The dashed line corresponds to a linear regression. The black bars correspond to standard deviation observed across simulations. (**H**) Comparison of the inferred C_normalised_ parameter with the noise parameter of the simulated noisy Thomas point pattern. Each dot is the average result of 50 simulations. The dashed line corresponds to a linear regression. The black bars correspond to standard deviation observed across simulations. **(I)** Spatial pattern of 9 cell types from the MERFISH Tonsild dataset. For each PCF-SiM parameter, 3 patterns are shown with an increasing parameter value from to to bottom.

In parallel, we tested 6 different families of function that had to **i**) be strictly positive, **ii**) converge to 1 for large distances and iii) with only a limited number of parameters are able to display a large number of shapes (Figure 1B). These 6 families consist in:

– The **power-law** model that has already been used to model pcf in astronomy [Snethlage et al., 2002]
– The **exponential**, **gamma** and **generalized gamma** models, all three being derived from the respective exponential, gamma and generalized gamma probability distribution functions. The generalized gamma model extends the gamma model, which itself generalizes the exponential model.
– The **sigmoid** model, derived from the commonly used sigmoid function in the context of Hill equation
– The **beta-prime** model, which is also derived from a probability distribution function and can be considered as a negative control.

When looking at function fitting quality across all four datasets (Figure 1C, D, E and F), we observed that the gamma, generalized gamma and sigmoid models fitted the best. In contrast the simpler power law and exponential models (two parameters only) had lower R^2^, suggesting that such simple models are not flexible enough to model the diversity of observed pcfs. Finally, the beta-prime model had the lowest R^2^ for three of the datasets, in agreement with its ‘negative control’ status.

While the gamma, generalized gamma, and sigmoid models produced fits of comparable quality, we chose to focus on the sigmoid model because its parameters are directly interpretable. The function used in the sigmoid model is defined as:

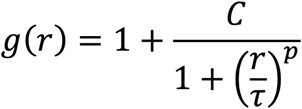

where **τ**, **p** and **C** are strictly positive parameters. This function is strictly decreasing, and reaches its half point when r = **τ** while **p** controls the steepness of the decline (Supplementary Figure 1D). Finally **C** is a scaling parameter (Supplementary Figure 1D) which after fitting is normalized with respect to **τ** and **p** (Methods) to obtain the C_normalised_ parameter.

We then decided to further investigate the interpretation of the fitted parameters. We first measured the ability of the sigmoid model to characterize the scale of cellular structures by simulating clusters of cells of various size through the generation of random Thomas cluster processes (Supplementary Figure 1E, Methods). We observed a near perfect correlation (R^2^=0.99) between the fitted **τ** parameter of the sigmoid model and the simulation cluster size parameter, suggesting that our approach is able to infer structure/ pattern characteristic size (Figure 1G). We performed a similar approach on a ring structure of varying width (Supplementary Figure 1F) and again observed a near perfect correlation between the fitted τ parameter and the structure size (Supplementary Figure 1G). Thus, we considered that the **τ** parameter of the sigmoid fit can be directly interpreted as a measure of the structure size.

We then wondered if the sigmoid model fit could also quantify the structure ‘neatness’, i.e the proportion of points or cells that are not localized in the structure and rather ‘noise’. We thus again simulated random Thomas cluster processes and added a variable proportion of randomly spatially distributed points (Methods). Interestingly, we observed that the noise parameter of this simulation (the ratio of noisy to clustered points) was perfectly correlating with the log transformed C_normalised_ parameter of the sigmoid model (Figure 1H), suggesting that the C_normalised_ parameter can be directly interpreted as a signal to noise parameter.

Finally we visually inspected the spatial distribution of the different cell types detected in the CosMX tonsil sample and observed that indeed the **τ** parameter value could be directly linked to the cellular structure size, while the the **p** and C_normalised_ parameters could be visually linked to the structure ‘sharpness’ and the proportion of ‘noisy cells’, i.e. randomly distributed cells, respectively (Figure 1I).

In sum, we developed a new approach, that we hereby call PCF-SiM (Pair Correlation Function – Sigmoid Modeling) that characterizes the spatial pattern of cells and can be directly and easily interpreted.

### Benchmark and robustness analysis of the sigmoid pcf fitting method

As several tools derived from the field of point pattern analysis have already been developed, we compared their outputs to those obtained with our approach. We first focused on the commonly used Clark–Evans (CE) aggregation index, which is defined as the ratio between the observed and expected mean nearest-neighbor distances. A CE index lower than 1 indicates spatial aggregation, whereas a value greater than 1 indicates a regular (dispersed) pattern. We computed the CE index for each cell type in each dataset and then evaluated its correlation with each of the sigmoid model parameters across datasets. To our surprise we did not observe any clear, strong and reproducible correlation across datasets for any of the sigmoid parameters (Figure 2A to D), suggesting that our approach and the CE index are not overlapping, likely because the CE index focuses on local and short-range structures in contrast to our method. We thus decided to focus on another spatial statistics, the negative-binomial overdispersion index [Bost et al. 2025] which recapitulates the spatial pattern more globally than the CE index. We indeed observed a strong and reproducible correlation between the C_normalised_ as well as the **p** parameters, and the overdispersion index (Figure 2E to H). A more in-depth analysis revealed that these correlations were statistically significant for three of the four datasets (Supplementary Figure 2A to 2H), hence validating the link between the overdispersion index and the two C_normalised_ and **p** parameters.

**Figure 2.**
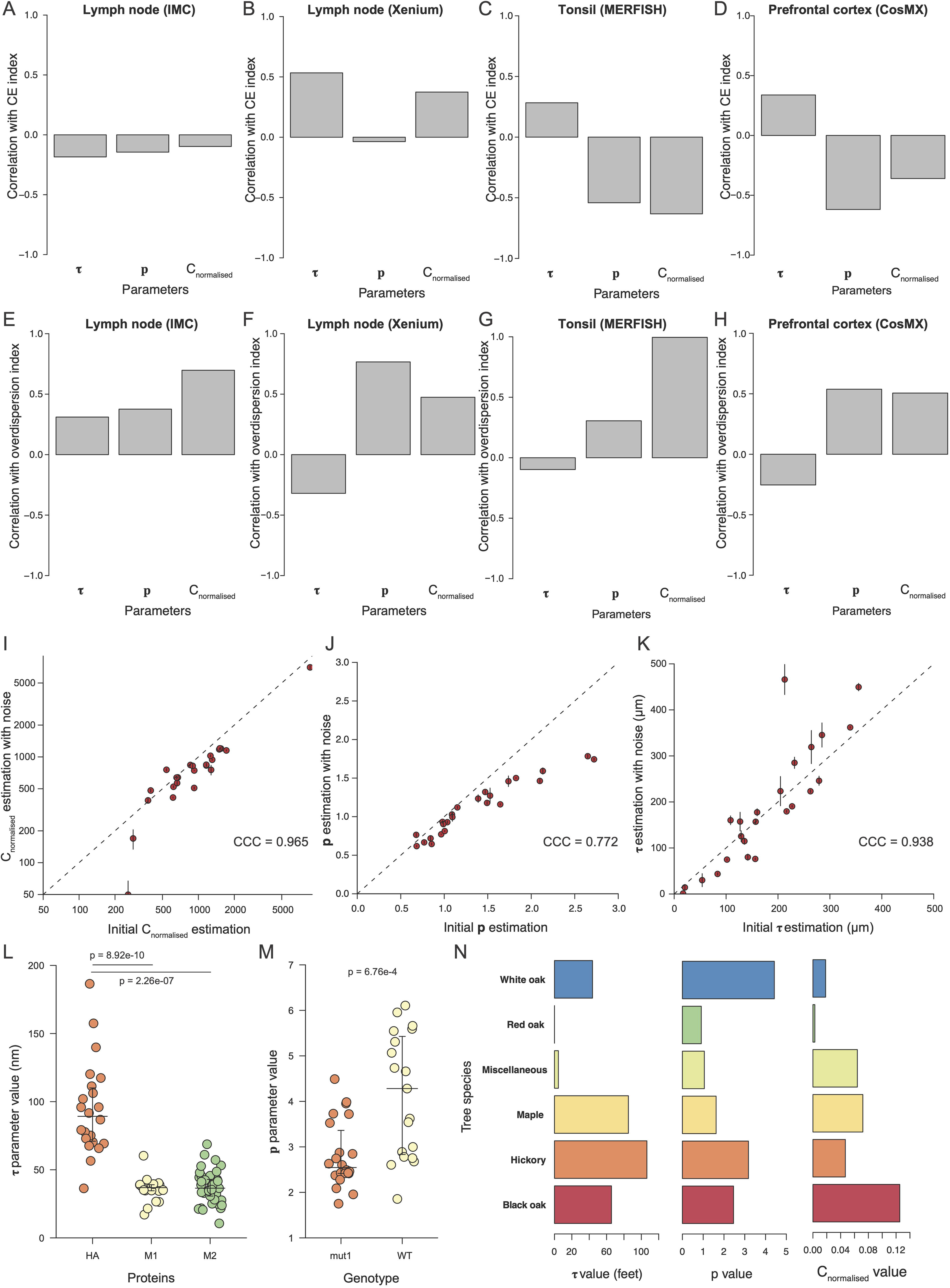
Benchmark and robustness analysis of the PCF-SiM framework. **(A)** Pearson’s correlation between the Clark-Evans index and the three PCF-SiM fitted parameters for the IMC Lymph node dataset. (**B**) Pearson’s correlation between the Clark-Evans index and the three PCF-SiM fitted parameters for the Xenium Lymph node dataset. (**C**) Pearson’s correlation between the Clark-Evans index and the three PCF-SiM fitted parameter for the MERFISH Tonsil dataset. (**D**) Pearson’s correlation between the Clark-Evans index and the three PCF-SiM fitted parameters for the CosMX Prefrontal cortex dataset. **(E)** Pearson’s correlation between the overdispersion index and the three PCF-SiM fitted parameters for the IMC Lymph node dataset. **(F)** Pearson’s correlation between the overdispersion index and the three PCF-SiM fitted parameters for the Xenium Lymph node dataset. **(G)** Pearson’s correlation between the overdispersion index and the three PCF-SiM fitted parameters for the MERFISH Tonsil dataset. **(H)** Pearson’s correlation between the overdispersion index and the three PCF-SiM fitted parameters for the MERFISH Tonsil dataset. **(I)** Comparison of the initially estimated C_normalised_ parameter and the estimated C_normalised_ parameter after label corruption. Each dot corresponds to a cell type and is the result of 100 simulations. Dashed line corresponds to the x=y line. The black bars correspond to the standard deviation observed across samples. **(J)** Comparison of the initially estimated **p** parameter and the estimated **p** parameter after label corruption. Each dot corresponds to a cell type and is the result of 100 simulations. Dashed line corresponds to the x=y line. The black bars correspond to the standard deviation observed across samples. **(K)** Comparison of the initially estimated **τ** parameter and the estimated **τ** parameter after label corruption. Each dot corresponds to a cell type and is the result of 100 simulations. Dashed line corresponds to the x=y line. The black bars correspond to the standard deviation observed across samples. (**L**) Estimated **τ** parameter for the different proteins of the Influenza data. The large horizontal lines correspond to the median while the two smaller horizontal bars correspond to the interquartile ranges. All “prism-like” plots in the manuscript are using the same graphical codes. P-values were computed using Tukey’s honestly significant difference test. (**M**) Estimated **p** parameter for the M2 protein according to the protein genotype. P-value was computed using a regular Welsh test. (**N**) Estimated PCF-SiM parameters for the different tree species from the Lansing Woods dataset.

We next evaluated the robustness of our method. To this end, we used the original cell annotations from the tonsil MERFISH dataset and, for each cell type, randomly swapped the locations of 20% of its cells with those of another cell type before performing a PCF-SiM analysis. This procedure introduced controlled label corruption while preserving the overall cell density distribution. We found that such perturbations only marginally affected the fitted values obtained from the PCF-SiM analysis (Figure 2I–K). Specifically, when assessing reproducibility using the Concordance Correlation Coefficient (CCC) [Lin, 1989], a metric that accounts for both correlation and scale differences unlike Pearson’s correlation, between the mean parameter values across samplings and the true values, we observed CCCs close to 1 for both the C_normalised_ and **τ** parameters (CCC=0.965 and 0.938 respectively). In contrast, the CCC was slightly lower, though still high, for the **p** parameter (0.772), and we noted a consistent underestimation of its values. This bias likely arises because label corruption produces less sharply defined cell clusters, resulting in lower **p** estimates. Overall, these results demonstrate that our method is highly robust to label corruption and, consequently, to imperfect cell annotations.

Finally, we sought to determine whether our approach could be applied beyond the context of biological tissue organization. To this end, we performed a PCF-SiM analysis on a publicly available electron microscopy dataset reporting the spatial locations of three influenza viral proteins, HA (hemagglutinin), M1, and M2, on cellular membrane sheets [Chen et al., 2008]. Our analysis revealed that the **τ** parameter associated with HA was significantly larger than those of M1 and M2 (Figure 2L), consistent with the known segregation of HA into lipid rafts, unlike the other two proteins (Supplementary Figure 2I and J). In addition, because this dataset included both wild-type and mutant forms of the M2 protein, we could assess the impact of the mutation on its spatial distribution. We observed a strong and significant decrease in the **p** parameter for the mutant M2 (Figure 2M, Supplementary Figure 2K and L), in agreement with the fact that the studied mutation impairs the protein’s ability to interact with other viral components and form well-defined spatial clusters required for viral budding [Chen et al., 2008]. Lastly, we applied our PCF-SiM framework to the Lansing Woods tree dataset, which records the locations and species classifications of trees within a 19.6 acre plot. Analysis of the resulting parameters revealed that white and red oaks exhibit an almost spatially random distribution, as indicated by their low C_normalised_ value parameter (Figure 2N). In contrast, hickory and maple trees display large, spatially coherent domains characterized by high τ values, observations consistent with the spatial patterns visible in the original dataset (Supplementary Figure 2M).

Altogether, the PCF-SiM framework is thus robust and can be applied to a diversity of contexts outside of biological tissue analysis.

### Characterization of the tissue structure alterations in a mouse colitis model

As our approach had so far been tested only on datasets consisting of individual tissue samples, we next examined whether it could be applied to a more complex dataset encompassing multiple samples and biological conditions. To this end, we reanalyzed a recently published, high-quality MERFISH dataset describing the cellular landscape of the mouse intestine during Dextran Sulfate Sodium (DSS)–induced colitis [Cadinu et al., 2024]. This dataset was particularly suitable due to its comprehensive temporal sampling (four time points) and the presence of several biological replicates for each condition. We began by examining the average **τ** and **p** parameter for each annotated cell cluster across all samples. Clusters displaying the highest **τ** values were B cells and macrophages (Figure 3A), followed by inflammation-associated fibroblasts (IAFs) and the smooth muscle cell cluster 1 (SMC1). Visual inspection confirmed these findings: B cell 1 clusters formed large, well-defined aggregates, likely corresponding to Peyer’s patches (Figure 3B, left panel), whereas IAF3 cells formed a diffuse ring-like structure (Figure 3B, right panel).

**Figure 3.**
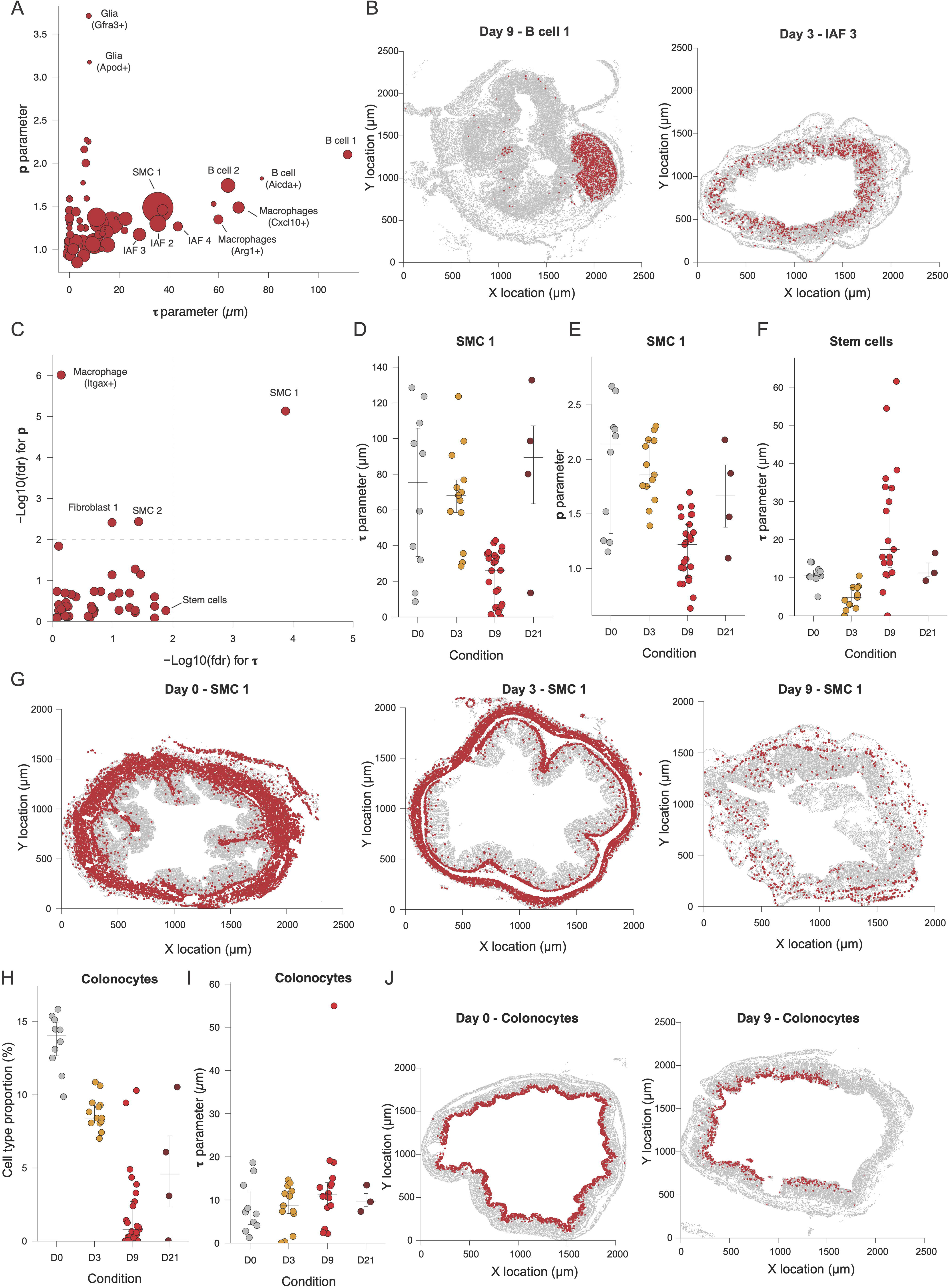
Characterization of mouse colon in the context of DSS-induced colitis. **(A)** Average **τ** and **p** parameters for each cell cluster. Each dot corresponds to a cell type and its size is proportional to the cell type abundance. **(B)** Spatial distribution of cells belonging to the Activated B cell 1 cluster (left) or the IAF3 cluster in representative samples. Cells of interest are colored in red while other cells are colored in grey. This color code will be conserved throughout the manuscript for similar plots. **(C)** Results of the differential structure analysis. The vertical and horizontal dashed lines correspond to the FDR thresholds of 1%. **(D)** Estimated **τ** parameter for the SMC 1 cluster across time points. **(E)** Estimated **p** parameter for the SMC 1 cluster across time points. **(F)** Estimated **τ** parameter for the stem cell cluster across time points. **(G)** Spatial distribution of cells belonging to the SMC 1 cluster on day 0 (left), 3 (middle) and 9 (right) in representative samples. **(H)** Proportion of colonocytes among imaged cells across time points. **(I)** Estimated **τ** parameter for the colonocyte cluster across time points. **(J)** Spatial distribution of cells belonging to the colonocyte cluster on day 0 (left) and 9 (right) in representative samples.

We then decided to perform a differential structure analysis, i.e. identifying which cell clusters had their **τ** and **p** parameters impacted through the course of the colitis. Interestingly, we observed that only one cell cluster, SMC 1, had both parameters affected in a statistically significantly manner (FDR < 0.0.1, Figure 3C) while three cell clusters (SMC 2, Fibroblast 1 and Macrophages Itgax+ clusters) had only their **p** parameter affected and none had only their **τ** parameter affected despite the stem cell cluster being borderline significant (Figure 3C). A more detailed look at the SMC 1 parameters kinetic revealed that indeed both the **τ** and **p** parameter values drastically dropped on day 9 post DSS treatment, before coming back around its initial values on day 21 (Figure 3D and 3E) while conversely, stem cells **τ** parameter was decreasing on day 3 and increasing on day 9 but in a highly heterogenous manner across samples (Figure 3F). Similarly, we observed that the SMC 2 **p** parameter was decreased from day 3 to day 21 while Fibroblast 1 **p** parameter was decreased on day 9 and 21 (Supplementary Figure 3A and 3B).

In order to validate our analysis, we inspected visually the slides to check that our analysis was not flawed: we indeed observed a massive thinning and disorganization of the ring formed by SMC 1 cells specifically on day 9 (Figure 3G). Similarly, we visually observed that the size of stem cell crypts was decreased on day 3 and increased on day 9 with respect to control, i.e. day 0 (Supplementary Figure 3C), as well as the loss of organisation of the SMC 2 cluster (Supplementary Figure 3D) on day 9. These results are in complete agreement with previous reports indicating a significant decrease of the muscularis propria thickness [Xu et al. 2020], as well as the expansion of stem cells upon acute injuries [Bohin et al. 2020] in the context of colon injury.

Finally, we wondered to which degree our structural analysis overlapped with a regular compositional analysis. We thus performed differential abundance analysis and realized that all cell clusters had their composition significantly affected by (FDR < 0.01), in contrast with the limited number of cell clusters displaying significant structural changes. To better understand this apparent paradox, we focused on the colonocyte cell cluster, a highly abundant cell cluster at homeostasis (>10% of the cells). While colonocyte abundance drops as soon as day 3, and falls to less than 5% on day 9 (Figure 3H), the fitted **τ** parameter was constant through time (Figure 3I), indicating that the structure size does not change. A visual inspection of the tissue section reveals that while in day 0 tissue sections the colonocytes form a thin yet uniform and complete ring (Figure 3J), this ring is incomplete in day 9 samples despite having the same thickness of ∼10µm, thus explaining the discrepancy between the structural and compositional analysis. In order to obtain more systematic results, we computed the correlation between each cell type abundance and each sigmoid model parameter: we observed an overall very limited correlation (Supplementary Figure 3E), especially for the **τ** parameter, which median correlation was close to 0. The PCF-SiM framework thus provides information orthogonal to the cellular composition.

Altogether, the PCF-SiM framework can be used to perform differential structure analysis and pinpoint at biologically relevant cell types, providing additional information to regular compositional analysis.

### Co-scaling analysis reveals structural interplay between cell types

So far, we have only applied the PCF-SiM framework to the point pattern associated to a single cell type. However, as every biological tissue encompasses a variety of cell types, describing spatial co-organisation between two cell types is essential to fully characterize tissue organisation. As a cross-pcf can be computed between two point patterns, i.e. between two cell types, we wondered whether we could apply our sigmoid model fitting approach to cross-pcf. We thus computed the cross-pcf between al cell types across every samples in the MERFISH colitis dataset before fitting the sigmoid models. To our surprised, we observed that in 24% of the cases, the sigmoid model could not be fitted while in 16% the model quality was low (R^2^<0.6), resulting in only 60% of correct fitting (Supplementary Figure 4A). In order to understand this surprisingly low success rate, we looked at the pcf correctly (Supplementary Figure 4B) and non-correctly fitted (Supplementary Figure 4C). We observed that while the correctly fitted cross-pcf had a shape similar to the regular pcf with decreasing values converging to 1, the non-correctly fitted pcf were strikingly different with non-monotonic behavior or values below 1, corresponding to repulsion between the two cell types at a given distance. As our sigmoid model can only be applied to strictly decreasing pcfs, this means that the PCF-SiM framework cannot be used to describe spatial co-organisation. Unfortunately, despite several attempts, we were not able to find a model which was able to fit to the entire diversity of cross-pcf while having a lo number of parameters.

As the direct modeling of the cross-pcf was not feasible, we realized that a different strategy could be used: Let two cell types A and B be part of the same biological structures, for instance forming respectively the inner and outer layers of a spheroid cellular aggregate (Figure 4A). Across tissue sections, the observed structure size might vary, but the ratio of the cell type A and B characteristic size (i.e. their tau parameter) should remain constant. Therefore we should observe a significant correlation between the **τ** parameter of the two cell types across samples. To test this strategy, we applied it to the MERFISH colitis dataset and in order to avoid spurious correlation caused by outliers, a common problem with Pearson’s correlation [Kim et al. 2015], we used the percentage bend correlation and its associated statistical test, a highly robust correlation measurement [Wilcox 1994]. Following multiple testing correction, we identified 10 significant co-scaling relationships (FDR < 0.01) spanning 10 different cell clusters (Figure 4B). Among them, several co-scaling relationship were between similar cell clusters, such as the relationship between the SMC 1 and 2, or the IAF 2 and IAF 3 clusters. A more in-depth analysis confirmed these results with a clear linear relationship between the **τ** parameters values in both cases (Supplementary Figure 4D and E), thus validating our initial co-scaling hypothesis. Indeed, in the case of the two SMC populations, we observe that they form two concentric rings with the SMC 1 and 2 respectively corresponding to the inner and outer layers of the muscularis propria (Supplementary Figure 4F).

**Figure 4.**
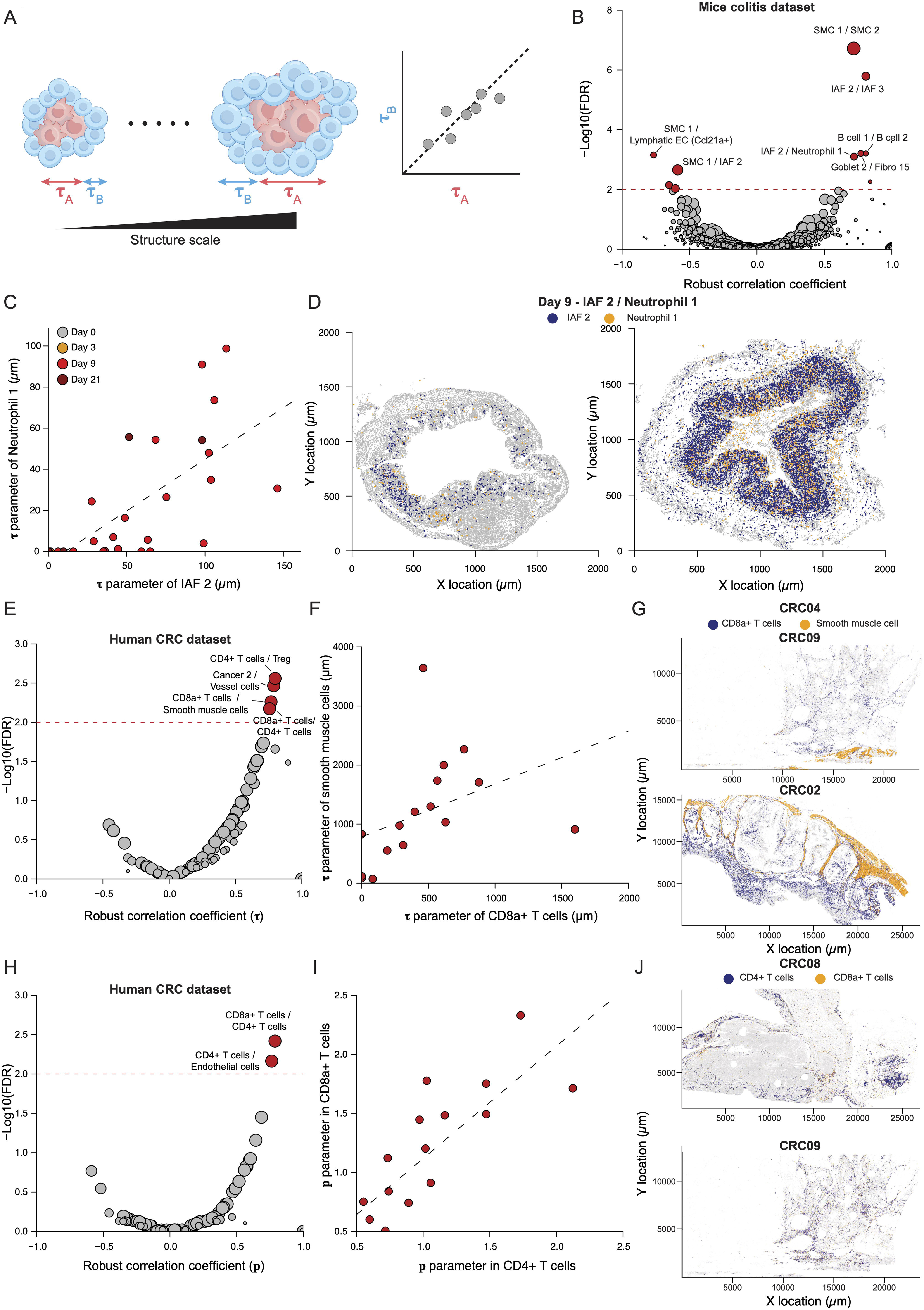
Validation of the co-scaling approach. **(A)** Illustration of the co-scaling approach**. (B)** Result of the co-scaling analysis for the mouse colitis dataset. Each dot corresponds to a cell cluster pair. Red dot correspond to signifiant co-scaling (FDR<0.01) while grey correspond to non-significant one. Dashed line corresponds to a significance threshold of 1%. Dot size is proportional to the number of samples used to compute the correlation. This graphical code will be used in similar plots throughout the manuscript. **(C)** Comparison of the **τ** parameter for IAF2 and Neutrophil clusters. Dashed line corresponds to a regular linear regression. **(D)** Spatial distribution of cells belonging to the IAF2 (blue) and Neutrophil 1 (orange) cluster in two day 9 samples. **(E)** Result of the co-scaling analysis for the CRC dataset. **(F)** Comparison of the **τ** parameter for CD8a+ T cells and smooth muscle cell clusters. Dashed line corresponds to a regular linear regression. **(G)** Spatial distribution of cells belonging to the CD8a+ T cells (blue) and smooth muscle cell (orange) clusters in two CRC samples. **(H)** Spatial distribution of cells belonging to the CD8a+ T cells (blue) and smooth muscle cell (orange) clusters in two CRC samples. **(I)** Result of the co-organisation analysis for the CRC dataset. **(I)** Comparison of the **p** parameter for CD8a+ T cells and smooth muscle cell clusters. Dashed line corresponds to a regular linear regression. **(J)** Spatial distribution of cells belonging to the CD4+ T cells (blue) and CD8a+ T cells (orange) clusters in two CRC samples.

We decided to further investigate heterotypic co-scaling relationships as they are more likely to be the results of cellular cross-talks and interactions. Among them, the Neutrophil 1 / IAF 2 pair is of particular interest as interactions between neutrophils and fibroblasts have been identified as driving various inflammatory diseases including Crohn’s disease [Gavriilidis et al. 2024]. We observed that this co-scaling was mostly driven by slides from day 9 mice (Figure 4C), in agreement with the fact that both cell types are recruited upon inflammation. Further visual inspection of the spatial data indicates that the two cell clusters tend to co-localize in the mucosa, suggesting that they likely interact, in agreement with the expression of known extracellular matrix remodeling genes (MMP8 and MMP9) by the Neutrophil 1 cluster [Cadinu et al., 2024]. As we had only tested our approach on a dataset containing a large number of samples derived from a well-controlled biological system, we wondered whether our approach could be applied to smaller and noisier datasets, typically datasets derived from small human cancer cohorts. We thus decided to re-analyze 16 samples from a cyclic immunofluorescence colorectal cancer (CRC) dataset [Lin et al. 2023]. Out of the 8761180 cells analyzed, we identified 19 cell clusters, encompassing 15 biologically meaningful cell types and states, including cancer, immune and stromal cell types (Supplementary Figure 4G). We applied our PCF-SiM framework to this dataset before running a co-scaling analysis. We identified 4 statistically significant co-scaling relationships, with the strongest being the CD4+ T cells and Tregs (Figure 4E). Similarly to the colitis dataset, we indeed observed clear linear relationships between the **τ** parameter values (Supplementary figure 4H to J, Figure 4F), albeit noisier than for the MERFISH colitis dataset. Interestingly, we observed a clear co-scaling between CD8a+ T cells and Smooth muscle cells (Figure 4G), in agreement with a recent clinical report showing a positive correlation between the number of tumor infiltrating lymphocytes and the skeletal muscle status in CRC patients [Daitoku et al. 2022].

Finally, we wondered if we could not only perform co-scaling analysis, but also co-organisation analysis, i.e. looking at the correlation between the **p** parameter values of two cell types. We thus computed the percentage bend correlation and its associated p-value for each cell cluster pair and observed that two cell cluster pairs (CD4+ T cells / CD8a+ T cells and CD4+ T cells / Endothelial cells) displayed a statistically significant correlation between their **p** parameter (Figure 4H). We decided to further investigate the CD4+ / CD8a+ pair, and indeed observed a very clear linear relationship between the **p** parameter values of both cell clusters (Figure 4I). Visual inspection of the spatial data indeed confirmed that in samples in which CD8a+ T cells form well-defined clusters, CD4+ T cells also form clearly defined cell clusters (Figure 4J, upper panel), while a spatially diffuse pattern of CD8a+ T cells will result in a diffuse pattern of CD4+ T (Figure 4J, lower panel), suggesting a shared regulation of CD4+ and CD8a+ T cells spatial aggregation.

In sum, we developed and validated a new approach to study how cell types are spatially co-organized in a tissue and validated it in-depth.

### Spatial structure characterization of Hashimoto’s Thyroiditis

Following the validation of our framework on a variety of publicly available datasets, we sought to apply it to complex and understudied pathological tissues. We focused on Hashimoto’s thyroiditis, an autoimmune disorder that is the most common cause of hypothyroidism in developed countries [Chaker et al., 2022]. The disease is characterized by the production of autoantibodies against the thyroid peroxidase (TPO) enzyme and by massive immune infiltration of the thyroid gland, ultimately leading to tissue damage and dysfunction [Klubo-Gwiezdzinska et al., 2022]. Although several genetic and environmental risk factors have been identified, the pathophysiology, and specifically the composition and spatial structure of the immune infiltrate, remains poorly understood.

To address this challenge, we used IMC to image four thyroid sections from patients with Hashimoto’s thyroiditis and four from healthy donors (Figure 5A). To optimize our experimental design, we constructed an antibody panel informed by a reanalysis of a publicly available scRNA-seq dataset [Hong et al. 2023] (Methods, Supplementary Table 1). We also implemented a stratified sampling strategy [Bost et al., 2025] to image stromal-cell–rich regions (“N regions”) separately from lymphocyte– and Tertiary Lymphoid Structure (TLS)–rich regions (“T regions”). The final dataset comprised 10 fields of view (FoVs) per patient sample, five N regions and five T regions, and 5 FoVs per healthy donor sample, capturing a broad diversity of tissue architectures and morphologies (Figure 5B).

**Figure 5.**
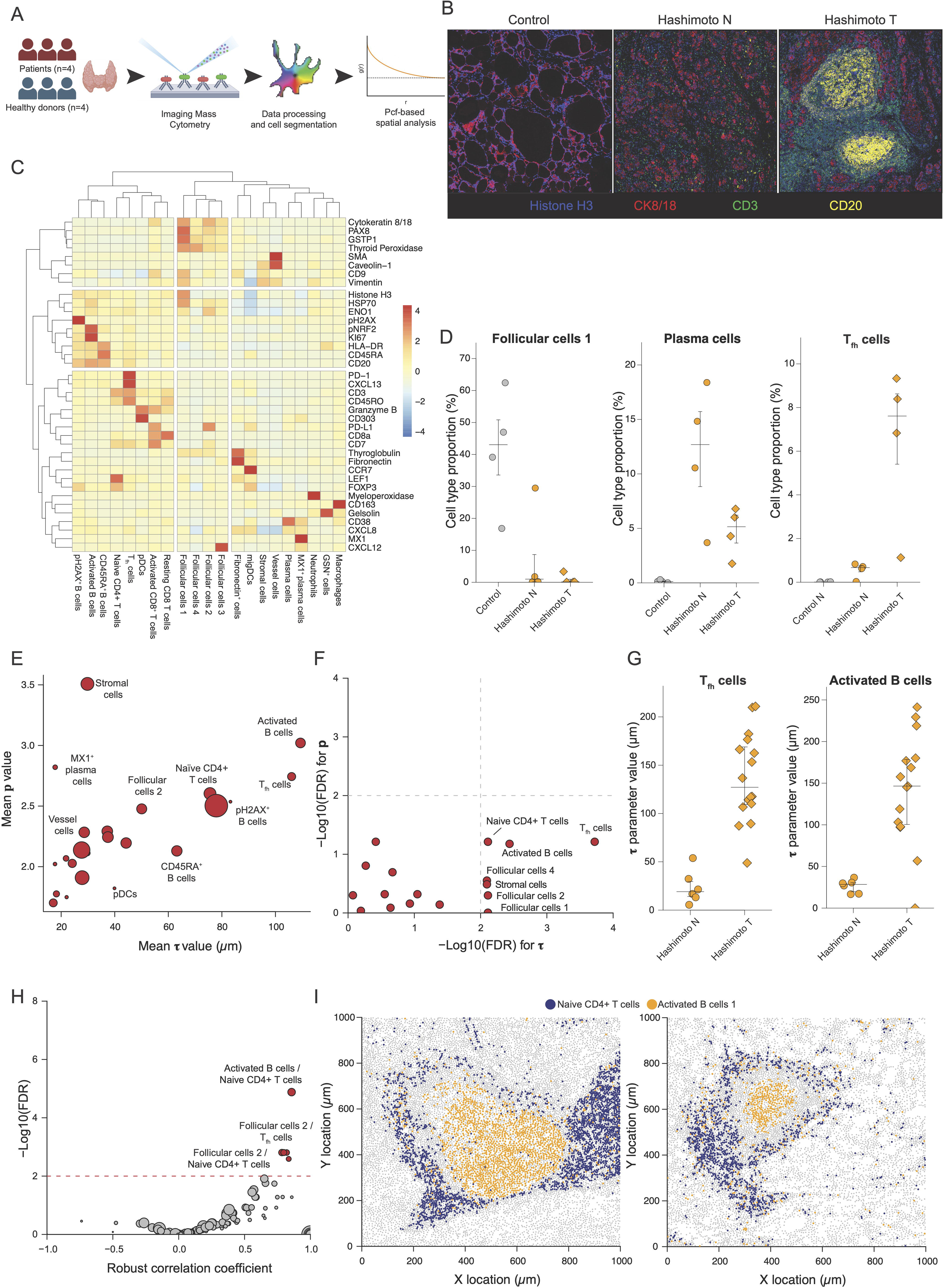
Characterization of Hashimoto’s thyroiditis associated tissue structure. (**A**) Experimental and computational framework for the analysis of Hashimoto’s thyroiditis. **(B)** Representative IMC images of the three types of FoV. All images are using the same color contrast. **(C)** Heatmap of the cluster gene gene expression profiles. The values are z-scaled by row. Associated hierarchical clusterings are performed using Ward’s criterion and euclidean distance. **(D)** Proportion of Follicular cells 1 (left), plasma cells (middle) and T_fh_ cells (right) across samples. **(E)** Average **τ** and **p** parameters for each cell cluster. Each dot corresponds to a cell type and its size is proportional to the cell type abundance. **(F)** Results of the differential structure analysis. The vertical and horizontal dashed lines correspond to the FDR thresholds of 1%. **(G)** Estimated **τ** parameter for the T_fh_ cell and Activated B cell 1 clusters across FoV types. **(H)** Result of the co-scaling analysis for the thyroiditis dataset. A threshold significance of 1% was used. **(I)** Spatial distribution of cells belonging to the naive CD4+ T cells (blue) and Activated B cells (orange) clusters in two representative Hashimoto T FoVs.

After cell segmentation and generation of a single-cell protein expression matrix, we performed unsupervised clustering (Methods) and identified 21 distinct cell clusters. These included four clusters corresponding to follicular cells, the hormone-producing epithelial cells of the thyroid, as well as diverse immune cell populations such as B cells, T cells, macrophages, and plasma cells (Figure 5C).

To characterize the compositional changes associated with disease, we performed Correspondence Analysis (CA) [Greenacre, 2007], a dimensionality reduction method analogous to PCA but designed for compositional data (Supplementary Figures 5A to C). The first CA dimension clearly separated FoVs from healthy and Hashimoto’s samples, with the separation driven by increased abundance of multiple immune cell types on the disease side and by stromal cell populations on the healthy side (Supplementary Figure 5C). Consistently, we observed a depletion of the homeostatic Follicular cells 1 cluster in Hashimoto’s samples (Figure 5D) alongside a marked recruitment of immune populations, including plasma cells and T follicular helper (T_fh_) cells (Figure 5D; Supplementary Figure 5D).

Having characterized disease-associated changes in tissue composition, we next investigated structural changes induced by the disease through our PCF-SiM framework. Immune cell clusters exhibited the largest **τ**, **p** and C_normalised_ values whereas the homeostatic follicular cell 1 cluster had the lowest C_normalised_ value, suggesting a highly structured immune infiltrate in contrast to the more randomly distributed stromal cells populations (Figure 5E, Supplementary Figure 5E). We then performed a differential structure analysis between the three types of FoVs and identified 6 cell clusters which had their **τ** parameter significantly varying between FoV types, while no cluster displayed significant differences in the **p** parameter (Figure 5F, FDR < 0.01). The most significant changes of the **τ** parameter were observed for T_fh_ cells and activated B cells, both of which showed an increase of the **τ** parameter in T regions compared to N regions of diseased samples (Figure 5G). Thus, although both populations are present in N and T regions, they form markedly larger and more coherent structures in T-region immune infiltrates.

To further probe higher-order tissue organization, we performed a co-scaling analysis. We identified five significant co-scaling relationships (Figure 5H), the most significant being the Activated B cells / Naive CD4+ T cells relationship, followed by the Follicular cells 2 / T_fh_ and Follicular cells 2 / Naive CD4+ T cells relationships. In all three cases, a clear linear co-scaling relationship was observed and driven predominantly by FoVs from disease samples (Supplementary Figure 5F to H), indicating that these co-scaling relationships are specific to the immune infiltrate. Regarding the Activated B cells / Naive CD4+ T cells relationship, visual inspection of the data revealed activated B cells form round aggregates surrounded by a shell of naive CD4⁺ T cells, with additional cell types interspersed between them, an organization characteristic of fully mature tertiary lymphoid structures (TLS) [Zhang et al., 2024] (Figure 5I). Finally, we investigated the two co-scaling relationships involving follicular cells 2, a subtype of follicular cells expressing the immune checkpoint PD-L1 (Figure 5C) and only found in patient samples (Supplementary Figure 5D). As PD-L1 is specifically induced by IFNγ, an inflammatory cytokine exclusively produced by T and NK cells [Boehmer et al. 2025], we hypothesized that the observed co-scaling is the result of the increased release of IFNγ in the context of large T cells aggregates, resulting in an increased induction of PD-L1 by neighboring follicular cells (Supplementary Figure 5I).

Altogether, by combining IMC with our PCF-SiM framework, we were able to characterize both compositional and structural alterations in thyroid tissue organization associated with Hashimoto’s thyroiditis.

### Analysis of uveal melanoma liver metastasis

Liver metastases, defined as tumors that have spread to the liver from a primary organ, are a major cause of cancer-related mortality, with more than 50% of cancer patients developing liver involvement during the course of their disease [Tsilimigras et al. 2021]. Several tumor types display a strong tropism for the liver, among which uveal melanoma (UM) is notable, with more than 90% of metastatic cases involving the liver. Liver metastasis is the leading cause of death in patients with UM, and median overall survival remains below two years despite the use of modern immunotherapeutic strategies [Carvajal et al. 2023], underscoring the urgent need for new therapeutic approaches.

In order to better characterize the tumor micro environment of UM liver metastasis, we imaged 8 tissues sections from UM patients with diagnosed liver metastasis (patients’ clinical features can be found in Supplementary Table 2) using the Xenium platform and a customized gene panel targeting both liver, melanocytes and immune genes (Supplementary Table 3) (Figure 6A). Each tissue section contained both large tumor and hepatic regions, thus allowing to study the interactions between the two tissues (Figure 6B). Following cell segmentation, we applied our TranspaceR pipeline to annotate the cells and identified 19 distinct clusters (Figure 6C). Among them we identified 7 cell clusters corresponding to cancer cells, as indicated by their expression of the melanocyte associated genes MLANA and TYRP1 (Supplementary Figure 6A). In addition to cancer cells, we identified several immune cell population such as Kupffer cells (MARCO+/CD163+), plasma cells (IGHM+), neutrophils (S100A9+) and CD8+ T cells (CD8+/GZMK+), as well as a variety of hepatic stromal cells ranging from healthy hepatocytes (SPP2+/F9+) to hepatic stellate cells (RGS5+/NOTCH3+) and fibroblasts (DCN+/ECM1+) (Figure 6C). In addition to these expected cellstypes, we also identified two unanticipated cell types: (**i**) inflamed hepatocytes characterized by their expression of the acute phase proteins SAA1 and SAA2 (Supplementary Figure 6B) and the interferon-inducible chemokine CXCL9 (Figure 6C) and (**ii**) lymphatic endothelial cells expressing the lymphocyte-attracting chemokine CCL19 and CCL21 (Figure 6C). While these chemokines can be produced in the tumor microenvironment either by lymphatic endothelial cells [Miao et al. 2021] or fibroblast reticular cells [Zhang et al. 2024], we considered that these cells were lymphatic endothelial cells due to their expression of the GATA2 [Linnemann et al. 2011] and LAMA4 [Stenzel et al. 2011] genes. Identified cell types strongly vary in terms of abundance, with healthy hepatocytes being on average the most common cell type, followed by Kupffer cells and CD8+ T cells (Figure 6D, left panel). In contrast, cancer cells were on average among the rarest cell types as they were highly patient specific, as validated by the significantly lower entropy of cancer cell type distribution across patients compared to non-cancer cell types (Figure 6D, right panel). Finally, both inflamed hepatocytes and lymphatic endothelial cells were among the rarest cell types, representing on average around 1% of the cells (0.99% and 1.08% respectively) but were detected in all patient samples (N>500), suggesting that these cell types are not patient specific and are a consistent feature of UM liver metastasis micro environment.

**Figure 6.**
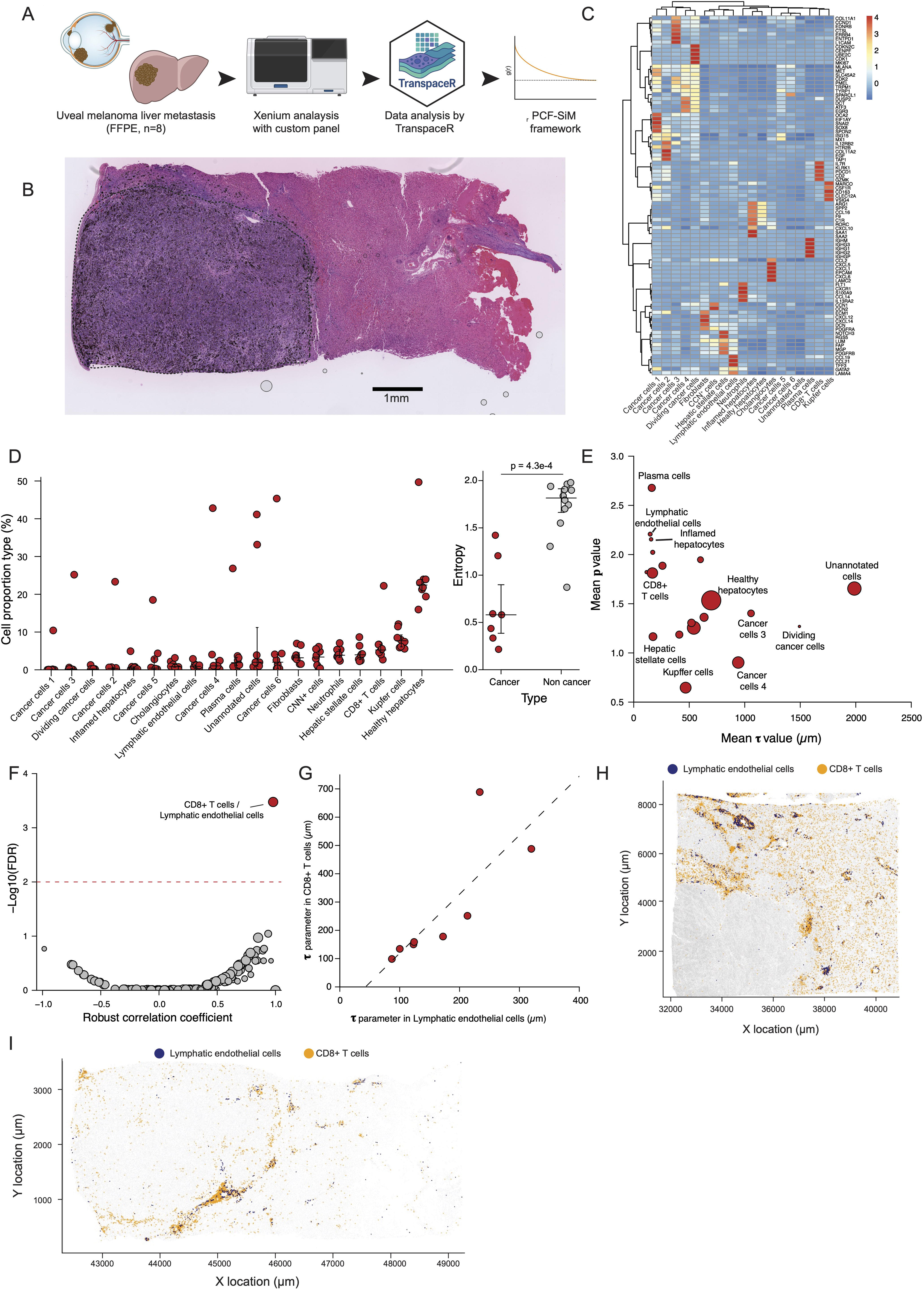
Dissection of uveal melanoma liver metastasis cellular structure. **(A)** Experimental and computational framework for the analysis of UM liver metastasis. **(B)** Whole-slide image of a hematoxylin and eosin stained of one of the patient samples. The dashed black curve delimitates the tumor. **(C)** Heatmap of the cluster gene gene expression profiles. The values are z-scaled by row. Associated hierarchical clusterings are performed using Ward’s criterion and euclidean distance. **(D)** Left panel: Cell type proportions across samples. Cell types are ordered based on their median abundance. Each dot corresponds to a sample. Right panel: Cell type entropy distribution of cancer and non cancer cell types. P-value was computed with a regular Welsh test. **(E)** Average **τ** and **p** parameters for each cell cluster. Each dot corresponds to a cell type and its size is proportional to the cell type abundance. **(F)** Results of the differential structure analysis. The vertical and horizontal dashed lines correspond to the FDR thresholds of 1%. **(G)** Estimated **τ** parameter for the CD8+ T cell and Lymphatic endothelial cells clusters across FoV types. **(H)** Spatial distribution of cells belonging to CD8+ T cells (blue) and CD8+ T cells (orange) clusters in a UM liver metastasis sample with the largest CD8+ T cell infiltrate (i.e. **τ** value). **(I)** Spatial distribution of cells belonging to CD8+ T cells (blue) and CD8+ T cells (orange) clusters in the UM liver metastasis sample with the lowest CD8+ T cell infiltrate (i.e. **τ** value).

We then applied our PCF-SiM framework and observed a strong variation of the fitted parameter values across cell types (Figure 6E): While some cancer cell types formed large structures with a **τ** parameter bigger than 1mm, corresponding to the large cancer masses (Figure 6B), more than half of the cell types had a **τ** parameter lower than 500µm, including lymphatic endothelial cells and inflamed hepatocytes. A similar variability was observed with the **p** parameter, with plasma cells, lymphatic endothelial cells and inflamed hepatocytes having the highest **p** parameter values, in contrast to Kuppfer cells. Visual inspection of the spatial data confirmed this analysis, with the plasma cells, lymphatic endothelial cells and inflamed hepatocytes forming small and well-defined structures clusters (Supplementary Figure 6C to E) while Kupffer cells formed poorly-defined cell aggregates (Supplementary Figure 6F).

Despite the low number of available samples, we performed a co-scaling analysis in order to detect possible interactions between cell types. We observed only one statistically significant co-scaling relationship (Figure 6F), corresponding to a clear and linear co-scaling between lymphatic endothelial cells and CD8+ T cells (Figure 6G). Visual inspection confirmed the result of our co-scaling analysis, with samples displaying clear lymphatic endothelial structures having clear and large CD8+ T cells aggregates (Figure 6H), while samples with nearly no clear endothelial structures exhibited limited CD8+ T cell infiltration (Figure 6I). As lymphatic endothelial cells have been described as key players of tumor immune surveillance [Karakousi et al. 2024], it is likely that in the context of UM liver metastasis, lymphatic endothelial cells are required for the effective recruitment of immune cells in the tumor microenvironment.

### Spatial structure characterization requires whole slide imaging and is not compatible with tumor micro arrays

Recent studies by us and others have illustrated the importance and complexity of experimental design for multiplexed imaging, especially due to the significant cost of such experiments [Bost et al. 2023], [Baker et al. 2023], [Bost et al. 2025]. However these pioneer studies only focused on the optimisation of the experimental design to maximize the number of detected cell types or to optimise the quality of the tissue composition estimation, and did provide any clear guidelines for optimising tissue structure characterization. In order to fill this gap we thus investigated the impact of FoV size on the output of our PCF-SiM framework: indeed we have so far only used whole-slide dataset, yet a large number of multiplexed imaging studies are based on Tumor Microarrays (TMA) [Camp et al. 2000], or on small FoV of ∼1mm^2^. We thus used the MERFISH tonsil dataset to simulate the acquisition of small FoVs of various sizes (0.5 to 2mm width), before applying our PCF-SiM framework to the subsampled data (Figure 7A). We first observed that due to the frequent absence of some cell types in these FoVs, no fit could be computed for a large number of samplings (Figure 7B). Indeed for the 500µm FoV, no fit could be computed in more than 50% of the sampling for more than half of the identified cell types. Although this fraction decreases with increasing FoV sizes, it remained significant even with 2000µm FoV. We also observed that fitting quality, measured by R^2^, was increasing with the FoV size and that for 500µm FoV, only a small fraction of cell types displayed on average a high R^2^ (R^2^>0.8) (Figure 7C). We then evaluated the quality of the PCF-SiM estimation: we observed that while the average estimated **τ** and **p** parameters would correlate with the real **τ** and **p** parameters values, for instance for 1000µm FoV (Figure 7D and 7E), the **τ** parameter would tend to be underestimated while the **p** parameter would be overestimated (Figure 7F and G). Interestingly, the quality of the estimation, measured by the CCC, increases with FoV size for both parameters (Figure 7H and I), suggesting a decrease in the bias for both parameters. In order to check the validity of our results, we applied the same approach to the Xenium Lymph node dataset, and observed similar results regarding the absence of fit (Supplementary Figure 7A), the fitting quality (Supplementary Figure 7B), and the estimated **τ** and **p** parameters (Supplementary Figure 7C to F).

**Figure 7.**
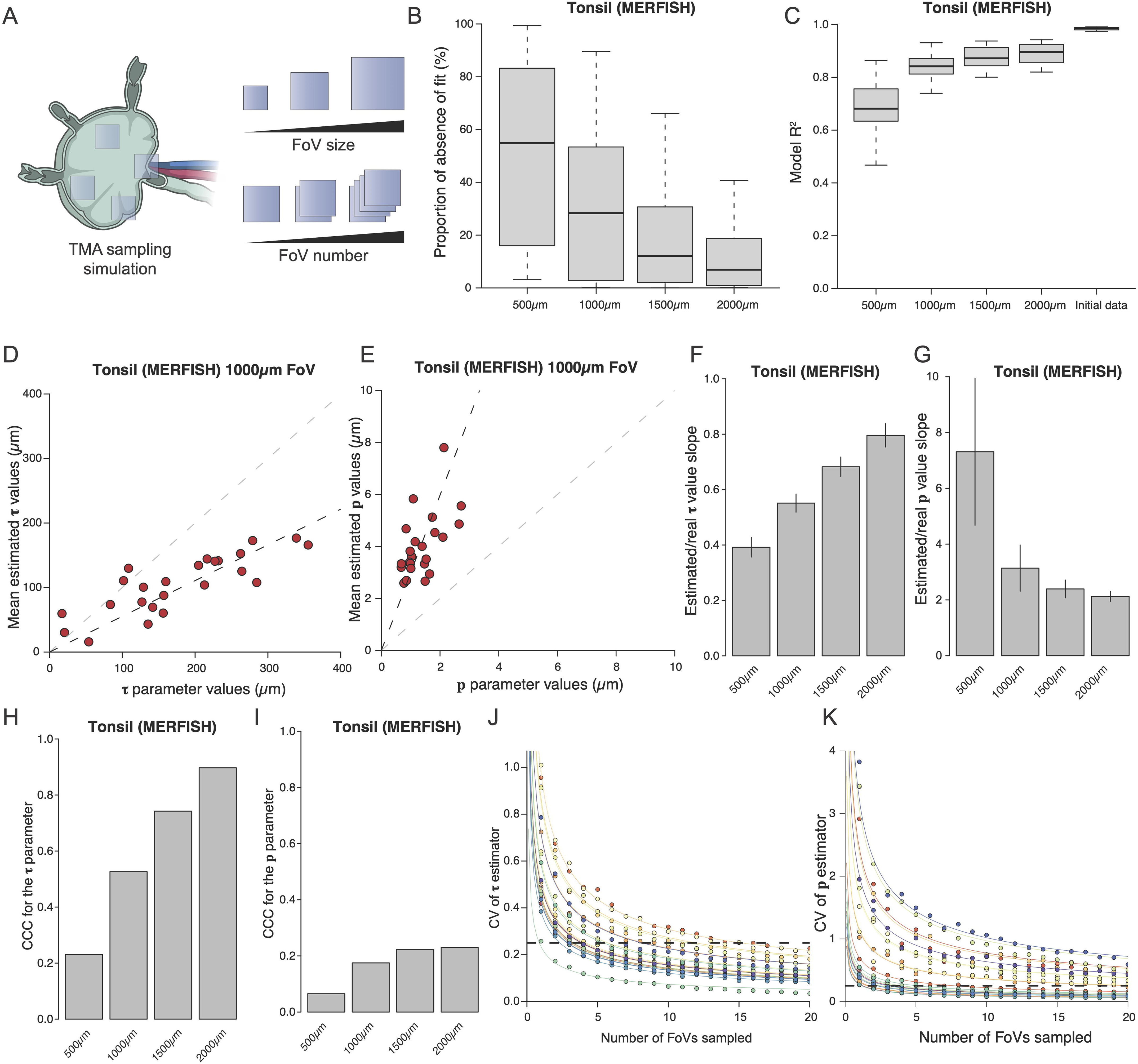
Effect of experimental design on the PCF-SiM framework. **(A)** Computational framework to simulate TMA acquisition. **(B)** Distribution of the proportion of absence of fit across cell types for different FoV size for the MERFISH tonsil dataset. The boxplot is based on N = 24 cell types. **(C)** Distribution of the R^2^ fit across cell types for different FoV size for the MERFISH tonsil dataset. The boxplot is based on N = 20 cell types. **(D)** Comparison of the initial **τ** parameter estimation and of the average **τ** estimate using 1000µm FoV on the MERFISH tonsil dataset. The black dashed line corresponds to the linear fit between the two variables. The grey dashed line corresponds to the line x=y. **(E)** Comparison of the initial **p** parameter estimation and of the average **p** estimate using 1000µm FoV on the MERFISH tonsil dataset. The black dashed line corresponds to the linear fit between the two variables. The grey dashed line corresponds to the line x=y. **(F)** Estimated slope between the **τ** parameter values and the average **τ** estimate across the sampled FoVs of different size. The vertical bars correspond to the slope estimate standard deviation. **(G)** Estimated slope between the **p** parameter values and the average **p** estimate across the sampled FoVs of different size. The vertical bars correspond to the slope estimate standard deviation. **(H)** CCC between between the **τ** parameter values and the average **τ** estimate across the sampled FoVs of different size. **(I)** CCC between between the **p** parameter values and the average **p** estimate across the sampled FoVs of different size. **(J)** Relationship between FoV number and the coefficient of variation of the **τ** estimator. Each color corresponds to a different cell cluster. The colored lines correspond to the inverse square root fit. The dashed line corresponds to the CV threshold of 25%. **(K)** Relationship between FoV number and the coefficient of variation of the **p** estimator. Each color corresponds to a different cell cluster. The colored lines correspond to the inverse square root fit. The dashed line corresponds to the CV threshold of 25%.

We finally wondered how many fields of views were necessary to obtain a non-noisy, yet biased estimate of the **τ** and **p** parameters. We thus computed the coefficient of variation of the mean estimate of the **τ** and **p** parameters for various FoV numbers. We observed that as expected, the coefficient of variation (CV) was proportional to the inverse square root of the number of sampled FoVs (Figure 7J and K), with highly variable initial CV across cell types. As we empirically considered that a CV of 25% had to be reached to obtain a reliable parameter estimation, this initial CV variation resulted in a variable number of required FoVs: for the **τ** parameter we observed that the minimal number of FoVs ranged from 2 to 16, with a median equal to 4 and an average close to 6 (5.95), while for the **p** parameter the 25% threshold could not be reached within 20 FoVs for 8 out of the 24 cell types (median of 3 and mean of 4.37 FoVs for the other cell types.) Altogether our sampling analysis shown that our framework produces biased estimates in the context of TMAs and small FoVs, and that in order to obtain non-noisy estimates, FoVs should be at least 1 mm^2^ while a minimum of 4 FoVs should be sampled, thus strongly limiting the usefulness of TMAs and small FoVs for tissue spatial characterization. Because our analyses were performed exclusively on lymphoid tissues, which are relatively disorganized and easier to sample than highly structured tissues such as tumors [Bost et al. 2023], this recommendation is likely conservative.

## Discussion

In the present study, we introduce a new framework, PCF-SiM, for characterizing the spatial distribution of cell types within tissues, designed to be both straightforward to implement and easily interpretable, even for non-specialists. We demonstrate the robustness and versatility of our approach across a wide range of publicly available datasets, as well as two clinically derived datasets generated in-house. Beyond describing spatial organization, our method enables differential structural analyses between groups of samples, providing a complementary perspective to traditional compositional comparisons. Furthermore, the co-scaling analysis reveals pairs of cell types that vary in a coordinated manner, indicating their participation in shared cellular structures and suggesting potential functional interactions within the tissue microenvironment.

We applied the PCF-SiM framework to a diverse collection of multiplexed imaging datasets spanning healthy and diseased tissues from both human and mouse samples, enabling a detailed characterization of their underlying spatial architecture. In a previously published colitis model [Cadinu et al. 2024], PCF-SiM revealed a profound disorganization of the muscularis propria and a marked expansion of stem cell crypts nine days after DSS induction, accompanied by a coordinated influx of specific neutrophil and fibroblast subtypes. When applied to a newly generated dataset of Hashimoto’s thyroiditis, the method uncovered a striking contrast between highly organized immune-rich regions and comparatively disordered stromal domains. Co-scaling analysis further suggested a functional interaction between infiltrating T cells and thyroid follicular cells, consistent with the emergence of PD-L1–expressing follicular cells in patient samples. Finally, analysis of a unique dataset of uveal melanoma liver metastasis demonstrated that several key cell types and states, including lymphatic endothelial cells and inflamed hepatocytes, form small but spatially coherent clusters at the tumor periphery. Despite the limited sample size (n = 8), our results additionally revealed that large lymphatic endothelial cell structures are strongly associated with substantial CD8⁺ T cell infiltration, pointing to a potential role of these cells in shaping local antitumor immunity.

While our approach represents a substantial methodological advance for the analysis of spatial cellular data, several extensions could further broaden its applicability and impact. In its current form, PCF-SiM relies on the pair correlation function as the foundation for all downstream analyses. Although powerful, this choice entails several assumptions about the underlying point pattern, the most important being isotropy. However, several biological spatial organizations are not isotropic, for instance keratinocytes in the skin form a largely planar layer, leading to strongly anisotropic spatial patterns. To address such cases, future developments could incorporate anisotropy into the model, in analogy with the extensions proposed for anisotropic variograms [Allard et al. 2016], thereby enabling more accurate characterization of tissues with directional or layered architecture. Furthermore, more sophisticated fitting strategies could be implemented, such as mixtures of sigmoid functions, to model more complex pcf behaviors, resulting from the presence of multiscale structures.

In addition to these methodological extensions, future work should explore the application of PCF-SiM to fully three-dimensional datasets. To date, we have evaluated our framework exclusively on 2D tissue sections, largely because most commercial multiplexed imaging platforms operate on thin slices rather than intact volumes. However, recent advances in tissue-clearing technologies and emerging approaches such as DNA microscopy [Qian et al. 2025] are making high-resolution 3D spatial transcriptomic and proteomic datasets increasingly accessible. As these volumetric data become more common, there will be a pressing need for analytical tools capable of capturing the added complexity of 3D spatial organization in a variety biological systems including human organ and organoids, underscoring the importance of extending our framework to three dimensions.

Finally, although we have applied our framework exclusively to multiplexed imaging datasets, rich in molecular information but typically limited in sample size, our approach could be extended to far larger cohorts. Given that our results indicate tissue microarrays (TMAs) are unsuitable for reliable structure analysis, an appealing direction would be to deploy PCF-SiM on digital pathology datasets, such as immunohistochemistry (IHC) or hematoxylin-and-eosin (H&E) slides annotated with modern deep-learning models [Jeong et al. 2025]. Because digital pathology collections often include thousands of patient samples, applying PCF-SiM at this scale would enable powerful statistical analyses, including survival modeling, thereby allowing structural tissue features to be evaluated as prognostic biomarkers. Such large-scale analyses could reveal clinically relevant spatial signatures that remain undetectable with conventional compositional or morphology-based approaches. Overall, the simplicity and robustness of our approach make it a powerful addition to the current toolbox of clinical pathologists.

## Methods

Computational analysis were performed using R version 4.03 (pcf fitting and analysis) and R version 4.5.1 (TranspaceR analysis). Python 3.8.5 was used for IMC image processing.

### Public spatial transcriptomic data download and processing

The three spatial transcriptomic datasets were obtained using the following procedure:

– We obtained the Xenium human lymph node dataset from the 10X Genomics website (https://www.10xgenomics.com/datasets/human-lymph-node-preview-data-xenium-human-multi-tissue-and-cancer-panel-1-standard). We reconstructed the metadata and expression files from the zarr files using a Python script from the TranspaceR pipeline [Mangane et al., 2025].
– We retrieved the MERFISH human tonsil dataset from the Gene Expression Omnibus platform using the accession number GSE282714. This dataset is derived from a recent publication [Zhao et al. 2025].
– We downloaded the gene count matrix and metadata for the human frontal cortex from the NanoString website (https://nanostring.com/products/cosmx-spatial-molecular-imager/ffpe-dataset/human-frontal-cortex-ffpe-dataset/).

We then used the TranspaceR pipeline to analyse and annotate these datasets. First we removed low-quality genes and cells. For the Xenium lymph node dataset parameters min_lib_size=10, max_lib_size=600, min_cell_radius=2, max_cell_radius=20 were used within the Curate_data() function. For for CosMx Human frontal cortex the parameters values were min_lib_size=10, max_lib_size=9500, min_cell_radius=2, max_cell_radius=15 whereas for the MERFISH Human tonsil these were min_lib_size=10, max_lib_size=2500, min_cell_radius=2, max_cell_radius=8. We calculated the variance with Excess_variance_ratio_NB(), zero proportion with Excess_zero_score_NB(), and Geary’s C scores with Geary_C_score() for each gene (as described in [Mangane et al. 2025]) using a significance p-value threshold of 0.01. We then used the expression of all the statistically significant genes for any score for cell clustering using the TranspaceR’s Cluster_makers() function with K parameter set to 50 for CosMx Human frontal cortex and Xenium Lymph node, and K = 100 for MERFISH Human tonsil. Finally we generated gene vs cluster heatmap using the Save_heatmap_markers() function by selecting the 5 markers of each cluster with the highest Log2 fold change compared to the overall gene mean expression. Cluster annotation was performed through literature curation and through the use of the Tabula Sapiens (https://tabula-sapiens.sf.czbiohub.org/) [THE TABULA SAPIENS CONSORTIUM].

### Pcf computation

For each cell type/cluster in each sample, the pcf was computed an in-house script based on the **spatstat** R package. First the pcf is only computed if the studied cell type has more than 50 points present in the sample of interest. Then a ppp object is build using the ppp() function from spatstat with default parameters, except for the window parameter. The point pattern window is defined as the smallest rectangle encompassing all cells and which is parallel/perpendicular to the x and y axis. As some cellular point patterns can contain up to hundred thousands of points, we did not directly computed the pcf but instead first computed the Ripley’s K function in highly optimized manner by Fast Fourrier Transform though the Kest.fft() function with the sigma parameter set to 1. The pcf is computed from the K function by derivation using the pcf.fv() function with the method parameter set to ‘b’, i.e. setting the pcf to 0 for r = 0, as two cells cannot physically overlap.

### Mathematical models for pcf fitting

We considered 6 different mathematical models to fit empirical pcf: the power-law, the exponential, the gamma, the generalized gamma, the sigmoid and the beta-prime models. These models were selected for the following reasons:

– The power law is already used in astronomy to model the pcf associated to galaxies distribution [Snethlage et al., 2002]
– The exponential, gamma and generalized gamma models are respectively derived from the exponential, gamma and generalized gamma probability distribution functions (pdfs). These pdfs are commonly used to model the distribution of positive random variable. While the exponential pdf is strictly decreasing, the gamma and generalized gamma pdfs are not necessarily monotonous, making them extremely flexible.
– The sigmoid model is derived from
– The beta-prime is derived from the eponymous pdf. This pdf models the distribution of a positive random variable and is characterized by its ‘heavy’ or ‘fat’ tail, i.e. its slow convergence to 0 [Bourguignon et al. 2020]. Therefore its ability to fit quickly converging pcf should be low and this model can serve as a negative control.

The functions and their associated parameters for each model are the following:

– The power-law model, where **τ** is the scale parameter and **α** the power exponent. Both parameters are strictly positive.

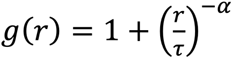
– The exponential model, where **τ** is the scale parameter and **C** the normalizing parameter. Both parameters are strictly positive.

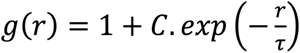
– The gamma model, where **τ** is the scale parameter, **α** a shape parameter**, C** the normalizing parameter. **τ** and **C** parameters are strictly positive while **α** is strictly bigger than –1.

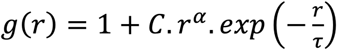
– The generalized gamma model, where **τ** is the scale parameter, **α** and **p** two shape parameters**, C** the normalizing parameter. **τ, p** and **C** parameters are strictly positive while **α** is strictly bigger than –1.

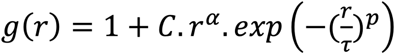
– The sigmoid model, where **τ** is the scale parameter, **p** a shape parameter and **C** the normalizing parameter. All parameters are strictly positive.

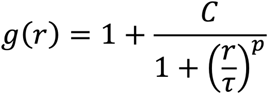
– The beta-prime model, where **α** and **β** are two shape parameters and **C** the normalizing parameter. All parameters are strictly positive.

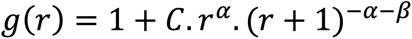

We empirically observed that the C parameter was not directly interpretable and has to be normalized with respect to the other parameters. We thus defined C_normalised_ as the normalized area under the curve of of g(r)-1.

### Fitting of the mathematical models

Each model was fitted by nonlinear least square using the nlsLM() function from the **minpack.lm** R package with the maximal number of iteration set to 1000. For each model, the following initial and lower bound values were used:

– For the power law model: initial **τ** and **α** values are respectively set to 100 and 2, lower bound values are both set to 0.
– For the exponential model: initial **τ** and **C** values are estimated by fitting a regular linear model where the log of the pcf is predicted as a function of distance using the lm() function. No lower bound values are used.
– For the gamma model: initial **τ**, **α** and **C** values are respectively set to 100, 2 and 100, lower bound values are respectively set 0, –0.999 and 0.
– For the generalized gamma model: initial **τ, α** and **p** values are respectively set to 10, –0.5 and 1, while the **C** parameter was to the maximal empirical pcf value observed. All lower bound values are set to 0 expect for **α** which was set to –0.999.
– For the sigmoid model: initial **τ** and **p** values are respectively set to 100 and 2, while the **C** parameter was to the maximal empirical pcf value observed. All lower bound values are set to 0.
– For the beta-prime model: initial **α**, **β** and **C** values were set to 15, 2 and 100. No lower bound values are used.

For all models, we computed the corresponding R^2^ as 1 minus the average ratio between the squared residuals and the pcf total variance. Residuals are extracted using the residuals() function. In each case, C_normalised_ was computed from the fitted parameters. In the case of the sigmoid model we have:

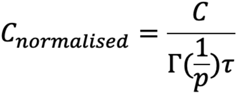

where Γ() is the gamma function.

### Point patterns simulations

We generated Thomas point pattern using the rThomas() function from the **spatstat.random** R package. For all simulations, the kappa parameter was set to 1, the mu parameter to 50, the and window to square of size 10. 50 point patterns were simulated for each scale parameter values. For the ring simulations, we simulated the distribution of 1000 points by first sampling a random angle between –π and π using the runif() function, before sampling random uniform value between –*w* and *w*, where *w* is half the radius thickness. This value is then added to 1 and multiplied by a radius of 2. These polar coordinates are then converted to cartesian coordinates and used to create a ppp object which associated window is a square of size 10. For each *w* value, 50 simulations were performed. For the simulation of noisy clusters, we first generated a Thomas point pattern as described above, before generating a Poisson point pattern using the rpoispp() function from the **spatstat.random** package, with the same window as the Thomas pattern. The lambda parameter was set to be the same as the Thomas pattern multiplied by a variable noise parameter. The two patterns were then superposed using the superimpose() function from the **spatstat.geom** package. The PcF-SiM framework was applied using the same method as previously.

### Comparison of PCF-SiM with other spatial metrics and robustness assessment

CE index was computed using the CE_interaction_tensor() function from the **Balagan** R package[Bost et al. 2023] with default parameter. The overdispersion index was computed by first computing quadrat counting using the quadratcount() function from the **spatstat.geom** package, with parameters nx and ny both set to 20. The obtain count data were used to fit Correlation was computed the base cor() function with the parameter use set to “pairwise.complete”. Regarding the label corruption step, the label corruption was performed individually for each cell type 100 times using the base sample() function. The concordance correlation coefficient was computed using an in-house function accordingly to the original publication [Lin, 1989].

### Application of PCF-SiM to non cellular spatial patterns

The influenza and Lansing Woods datasets were loaded from the **spatstat.data** R package using respectively the flu and lansing objects respectively. The PCF-SiM framework was applied similarly to other datasets.

### Analysis of the MERFISH colitis dataset

The cell_properties.csv file was downloaded from the data Dryad website (https://doi.org/10.5061/dryad.rjdfn2zh3). We used the ‘Tier2’ clustering level as an input for our pcf analysis. Differential structure analysis were performed using a regular ANOVA test with the aov() and anova() functions from the **stats** R package. Differential abundance analysis was performed through the fitting of Poisson generalized linear model (GLM): Briefly a Poisson GLM with a logarithmic link function was fitted for each cell type using the total number of cells and the time point as covariates. The significance of the time covariate was then assessed by a likelihood ratio test. The functions glm() with the parameter family set to “poisson” and anova() with the parameter test set to ‘LRT’ were used to do each step respectively. For both structure and compositional analysis, p-values were corrected for multiple testing using the p.adjust() function with the parameter method set to “fdr”.

### Cross-pcf computation and fitting

Cross-pcf were computed similarly to regular pcf, with the exception we added the cell type as a mark during the ppp object creation and used the Kcross() function instead of the Kest.fft() function. In order to improve obtain high-quality cross-pcf, we set the minimal number of points to 100 for each cell type. The sigmoid model was fitted similarly to regular pcf.

### Co-scaling analysis

As regular Pearson’s correlation is notoriously sensitive to outlier, we used the percentage bend correlation [Wilcox 1994] to study correlation between the **τ** parameter values of two cell types. We used the pball() function from the **WRS2** R package [Mair et al. 2020] with default parameters, i.e. a bending **β** parameter set to 0.2 and 500 random permutations used for p-value computation. The obtained p-values are corrected using the Benjamini-Hochberg procedure implemented in the p.adjust() function with the method parameter set to ‘fdr’.

### Analysis of the CRC dataset

Raw single-cell data from the CRC study [Lin et al. 2023] corresponding to samples CRC02 to CRC017 were downloaded from the corresponding Zenodo repository (https://zenodo.org/records/7554924). As the dataset was too large to be clustered and annotated directly, we randomly subsampled 5% of the cells before performing the clustering itself. We first removed the LaminABC and CDX2 channels from the dataset as those two features displayed an abnormally low variance to mean ratio. We then standardized each gene expression vector such that its variance was equal to 1 before performing singular value decomposition with the irlba() function from the **irlba** package with the nv parameter set to 20 (20 singular values computed). We then computed the associated K-nearest-neighbor (KNN) graph with the Knn() function from the **N2R** package using the latent dimensions as an input and by setting the k parameter to 30 and the indexType parameter to “L2”. Cell communities were then detected using the cluster_louvain() function from the **igraph** R package.

### Design of the IMC thyroid antibody panel

Raw scRNA-seq data from healthy human thyroid samples from [Hong et al. 2023] were downloaded from the Gene Expression Omnibus platform, using the GSE182416 accession number. The GSE182416_Thyroid_normal_7samples_54726cells_raw_count.txt.gz file was downloaded and loaded into R using the readRDS() base R function. Genes with less than 100 UMIs detected were removed. Highly variable genes were detected by performing a loess regression between the proportion of cells with non detectable expression and the average gene expression using the loess() function with default parameters. The 2000 genes with the highest positive residual values were used for further analysis. Data normalisation and clustering were performed using the Pagoda2 R package (https://github.com/kharchenkolab/pagoda2). The adjustVariance() function was run with parameter gam.k set to 5, the function calculatePcaReduction() with the nv parameter set to 50, the makeKnnGraph() function with k set to 30, type set to PCA and distance set to ‘cosine’. Finally the getKnnClusters() function was run with the method parameter set to multilevel.community. The obtained clustering was then used as an input for the Python package **scGeneFit** [Dumitrascu et al. 2021]. The function from this package was used with redundancy set to 0.25, method set to ‘pairwise’ and 40 markers. The initial marker list was then manually optimized with regards to available high-quality antibodies. Finally, all antibodies were validated using thyroid sections from healthy donors and patients.

### Thyroid section processing and IMC data acquisition

Thyroid samples in the form of FFPE blocks were obtained from the University Hospital Zurich (BASEC 2021-00417) and 5-µm thick sections were collected similarly to the melanoma samples and kept at –20°C until staining. Sections were dewaxed, rehydrated, and subjected to a heat-induced epitope retrieval step for 30 min at 95 °C in 10 mM Tris, pH 9.2, 1 mM EDTA. The sections were then incubated in blocking buffer (3% bovine serum albumin in Tris-buffered saline (TBS) with 0.1% Triton X-100) for 1 h at room temperature before incubation with a 38-antibody panel (Supplementary Table S1) diluted in blocking buffer overnight at 4 °C. In addition to the lanthanide-bound antibody, the antibody mix also contained two fluorescently labelled antibodies: anti pan-Cytokeratin antibody (clone AE1/AE3, 1.5µg/mL) labelled with the AF488 fluororophore, and an anti CD45 antibody (clone 2B11 & PD7/26, 2µg/mL) conjugated to the AF555 fluorophore. Nuclear staining was then performed by adding an iridium solution (5 nM) diluted in TBS (1:100 dilution) to the sample and incubating for 5 min. The samples were then washed three times (10 minutes per wash) in TBS and dried. Before IMC acquisition, the slides were imaged with with a slide scanner (Zeiss Axio Scan.Z1) at 20X magnification. Images were acquired using a Hyperion XTi Imaging System with the ablation frequency set to 800 Hz and the ablation energy set to 6 dB with X and Y steps set to 1 µm.

### Thyroid data pre-processing and analysis

The raw .mcd files generated by the Hyperion XTi Imaging System were converted into .tiff files using the **imctools** Python package (https://github.com/BodenmillerGroup/imctools) and more precisely using the McdParser and ImcWriter functions with default parameters. The obtained tiff files were then used as an input for cell segmentation using Cellpose version 2.2 [Stringer et al. 2021]. The Iridium channel was used as the nuclear channel, while the CD45RA, CD45RO, Cytokeratin 8/18 and HLA-DR channels were summed to generate the cytoplasmic channel. The model parameter was set to “tissuenet”, the diameter parameter set to 5, the flow_threshold set to 1, cellprob_threshold to –6, segmentation_type to ‘whole-cell’ and average_models set to True. Cell location and average signal intensity for each channel was computed using the skimage.measure.regionprops_table() function from the **skimage** Python package.

### Uveal melanoma liver metastasis section processing and Xenium data acquisition

All uveal melanoma patients were treated at Institut Curie, Paris, France, and signed informed consent. The project was approved by the ethics committee of Institut Curie (DATA190128).

Eight formalin-fixed paraffin-embedded (FFPE) uveal melanoma liver metastases were analyzed using 10X Genomics Xenium spatial transcriptomics. Samples were imaged with the immune-oncology panel (380 genes) to which a custom add-on probe sets against 100 additional genes was added. Probes were designed using the 10X Genomics online Xenium Panel Designer. The total list of genes is available in Supplementary Table 3. For each sample, a 5 µm section was collected, floated in a 37 °C water bath, and adhered to Xenium slides (10x, PN 1000460). Samples were baked at 42 °C for 3 h and stored in a desiccator at room temperature until processing. Samples were processed according to manufacturer protocols “FFPE_TissuePreparation, user guide CG000578 “GeneExp_CellSegmentation, User Guide CG0000749” and “Decoding & Imaging, User Guide CG000584”. Slides were stained with the Xenium sample prep reagents kit according to manufacturer instructions (10X, PN 1000460). Briefly, padlock probes were incubated overnight before ligation and rolling circle amplification. Subsequently, native protein autofluorescence was reduced with a chemical autofluorescence quencher. Slides were processed on a 10 Xenium Analyzer (software version 3.2), with ROIs selected to cover the different samples. Data was processed on the analyser through the standard 10X workflow to generate cell by gene and transcript by location matrices. Cell segmentation was performed using the multimodal segmentation kit (10X, PN 100662). This kit combines a DAPI nuclear staining, antibodies for ATP1A1, E-Cadherin, and CD45 for cell membrane boundaries, 18S Ribosomal RNA to label the cytoplasm, and alphaSMA/Vimentin antibodies for interior protein staining.

### Uveal melanoma liver metastasis data pre-processing and analysis

For the Uveal melanoma dataset, we aggregated the two Xenium objects from two different slides, each containing four tissue samples. Using our script for processing Xenium zarr data, we created a count matrix and metadata for each slide. We then combined the two datasets by stacking them and adjusted the cell locations on one slide. Next, we used the TranspaceR pipeline to analyze the combined dataset. We set specific parameters in the Curate_data() function: min_lib_size of 10, max_lib_size of 2500, min_cell_radius of 2, and max_cell_radius of 15. We calculated the variance score for each gene using the Excess_variance_ratio_NB() function and the zero proportion score with Excess_zero_score_NB(), applying a significance threshold of 0.01. For Geary’s C score, we used the Geary_C_score_multiple() function with a p-value threshold of 0.01 to analyze each sample separately and combined all significant genes across samples. We then performed cell clustering using the expression of the statistically significant genes from all these scores with the Cluster_makers() function from TranspaceR, setting the K parameter to 100. Finally, we created a heatmap showing the top 10 marker genes for each cluster using the Save_heatmap_markers() function.

### Sampling analysis

For both datasets on which the sampling was performed, we first sampled 500 square FoVs of various size (width of 500, 1000, 1500 and 2000µm) using the Random_spatial_sampling() function from the **Balagan** R package. If a FoV had less than 1000 cells it was removed from analysis in order to remove empty FoVs. The PCF-SiM framework was then applied to each FoV for each cell type and the obtained fit stored. The linear fit were performed using the lm() function with default parameters but without a null intercept. The standard deviation of the The concordance correlation coefficient was computed using an in-house function accordingly to the original publication [Lin, 1989]. For the computation of the estimator coefficient of variation for **τ** and **p** we used the following approach: for a given number of FoVs *n*, we randomly sample *n* estimates of the parameter for a given FoV size before averaging the sampled estimations. This step is repeated 500 times and the obtained values used to compute the standard deviation of the estimator with the function sd(). The coefficient of variation was obtained by dividing the standard deviation by mean estimated parameter. The relationship between the coefficient of variation and the number of fit was computed using the lm() function with no intercept.

## Data and code availability

All scripts used in this paper are accessible in a GitHub repository (https://github.com/BOSTLAB/PCFSiM-Paper). We created an R package, **PCFSiM**, freely accessible on GitHub (https://github.com/BOSTLAB/PCFSiM/). All annotated datasets are freely accessible on a Zenodo repository (https://zenodo.org/records/17867607). All raw data generated for this manuscript will be uploaded upon publication of the manuscript.

## Legends

**Supplementary Figure 1.**
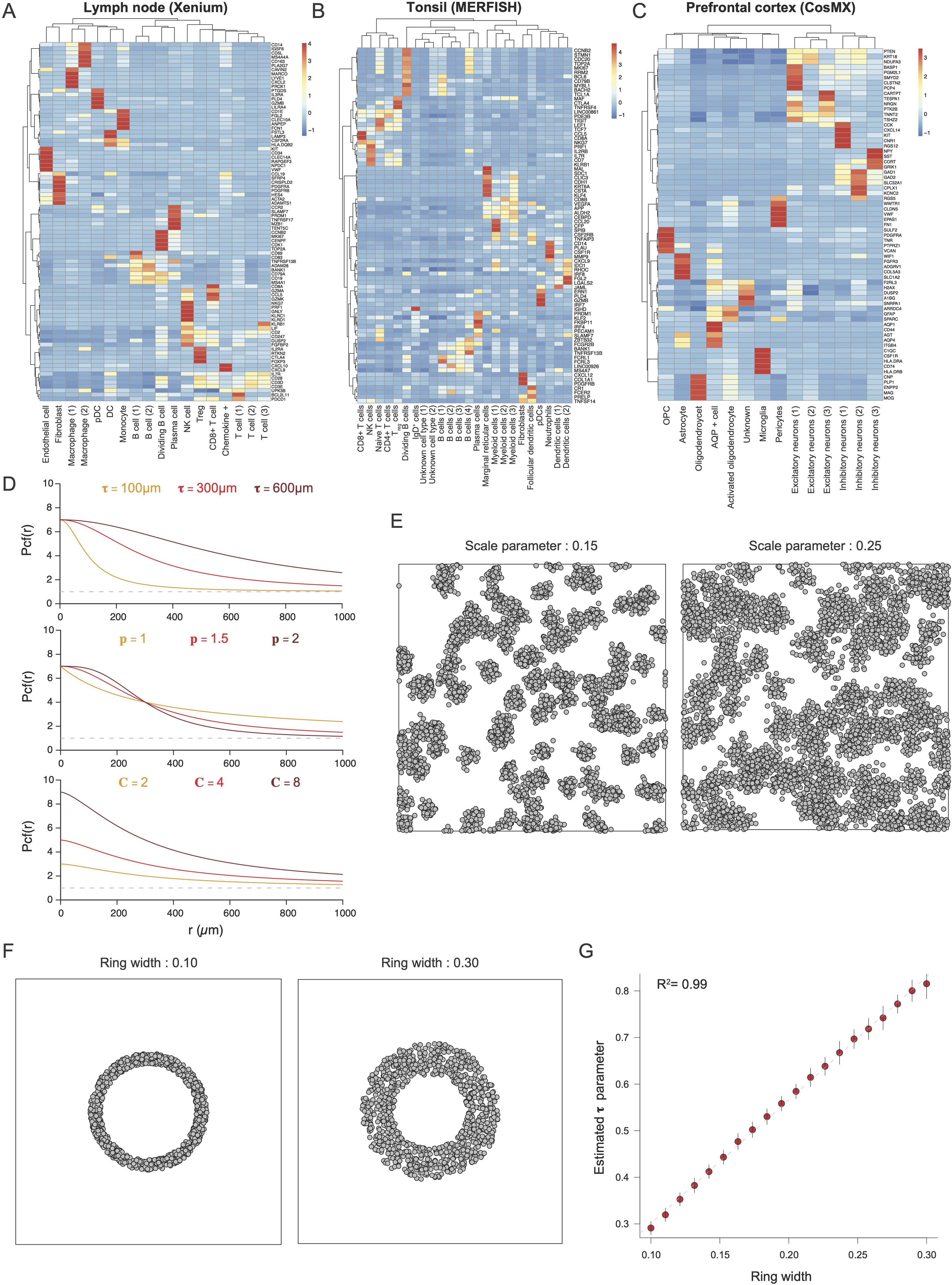
(**A**) Heatmap of the cluster gene gene expression profiles for the Xenium Lymph node dataset. Values are z-scaled by row. **(B)** Heatmap of the cluster gene gene expression profiles for the MERFISH Tonsil dataset. Values are z-scaled by row. **(C)** Heatmap of the cluster gene gene expression profiles for the CosMX Prefrontal cortex dataset. Values are z-scaled by row. **(D)**. Impact of the **τ** (top panel), **p** (middle panel) and **C** (bottom panel) parameters on the sigmoid function. **(E)** Representative simulated Thomas clustered point patterns with two different scale parameter values. **(F)** Representative simulated ring point patterns with two different ring thickness parameter values. **(G)** Comparison of the inferred **τ** parameter with the scale parameter of the simulated Thomas point pattern. Each dot is the average result of 50 simulations. The dashed line corresponds to a linear regression. The black bars correspond to standard deviation observed across simulations.

**Supplementary Figure 2.**
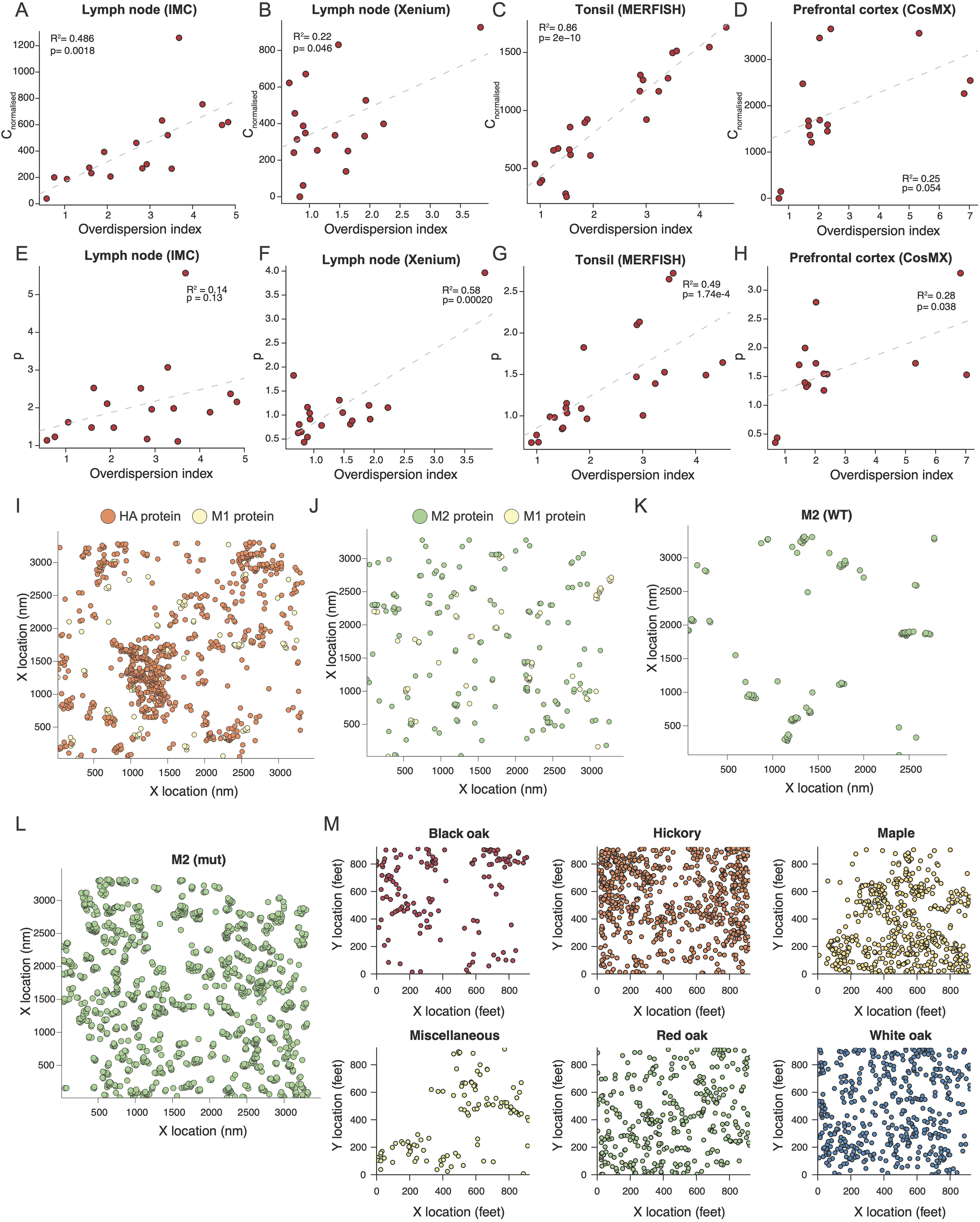
(**A**) Scatter plot of the overdispersion index and of the C_normalised_ parameter for the IMC lymph node dataset. The dashed line corresponds to a regular linear regression. **(B)** Scatter plot of the overdispersion index and of the C_normalised_ parameter for the Xenium lymph node dataset. The dashed line corresponds to a regular linear regression. **(C)** Scatter plot of the overdispersion index and of the C_normalised_ parameter for the MERFISH tonsil dataset. The dashed line corresponds to a regular linear regression. **(D)** Scatter plot of the overdispersion index and of the C_normalised_ parameter for the CosMX Prefrontal cortex dataset. The dashed line corresponds to a regular linear regression. **(E)** Scatter plot of the overdispersion index and of the **p** parameter for the IMC lymph node dataset. The dashed line corresponds to a regular linear regression. **(F)** Scatter plot of the overdispersion index and of the **p** parameter for the Xenium lymph node dataset. The dashed line corresponds to a regular linear regression. **(G)** Scatter plot of the overdispersion index and of the **p** parameter for the MERFISH tonsil dataset. The dashed line corresponds to a regular linear regression. **(H)** Scatter plot of the overdispersion index and of the **p** parameter for the CosMX Prefrontal cortex dataset. The dashed line corresponds to a regular linear regression. **(I)** Spatial distribution of HA and M1 influenza proteins in a representative electron microscopy sample. **(J)** Spatial distribution of M1 and M2 influenza proteins in a representative electron microscopy sample. **(K)** Spatial distribution of the M1 protein in a representative electron microscopy sample with the wild type M1 protein. **(L)** Spatial distribution of the M1 protein in a representative electron microscopy sample with the mutated M1 protein. **(M)** Spatial patterns present in the Lansing Woods tree dataset.

**Supplementary Figure 3.**
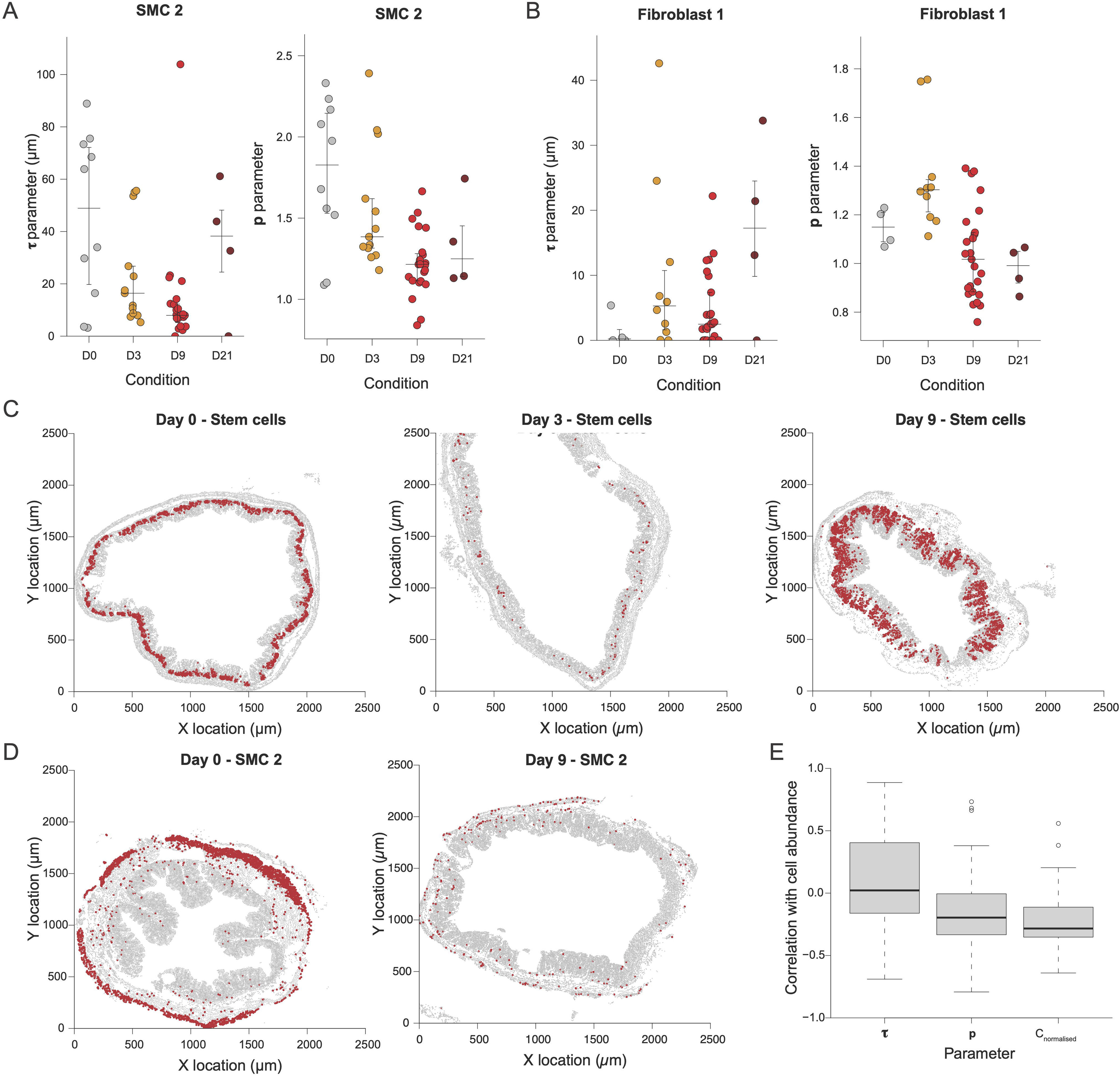
(**A**) Estimated **τ** (left) and **p** (right) parameters for the SMC 2 cluster across time points. **(B)** Estimated **τ** (left) and **p** (right) parameters for the Fibroblast 1 cluster across time points. **(C)** Spatial distribution of cells belonging to the Stem cell cluster on day 0 (left), 3 (middle) and 9 (right) in representative samples. **(D)** Spatial distribution of cells belonging to the SMC 2 cluster on day 0 (left) and 9 (right) in representative samples.

**Supplementary Figure 4.**
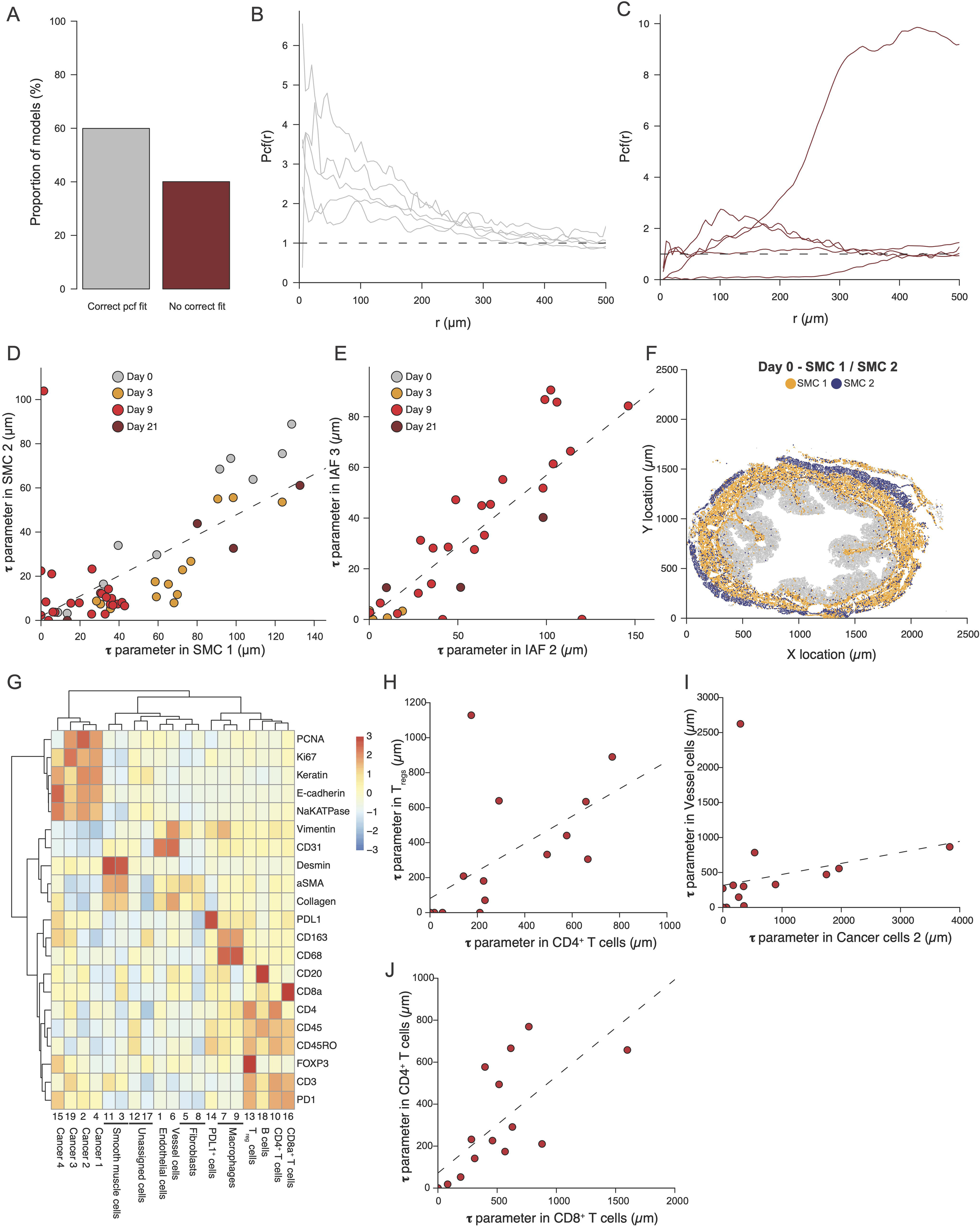
(**A**) Proportion of cross-pcf model that displayed a correct or a non correct fit. **(B)** Representative cross-pcf to which the sigmoid model fitted correctly. **(C)** Representative cross-pcf to which the sigmoid model **<u>did not</u>** fit correctly. **(D)** Comparison of the **τ** parameter for SMC 1 and SMC 2 clusters. Dashed line corresponds to a regular linear regression. **(E)** Comparison of the **τ** parameter for IAF 2 and IAF 3 clusters. Dashed line corresponds to a regular linear regression. **(F)** Spatial distribution of cells belonging to the SMC 2 (blue) and SMC 1 (orange) cluster a representative day 0 sample. **(G)** Heatmap of the cluster gene gene expression profiles for the CRC dataset. Values are z-scaled by row. **(H)** Comparison of the **τ** parameter for CD4+ T cells and Tregs. Dashed line corresponds to a regular linear regression. **(I)** Comparison of the **τ** parameter for Vessel cells and Cancer cells 2. Dashed line corresponds to a regular linear regression. **(J)** Comparison of the **τ** parameter for CD4+ T cells and CD8+ T cells. Dashed line corresponds to a regular linear regression

**Supplementary Figure 5.**
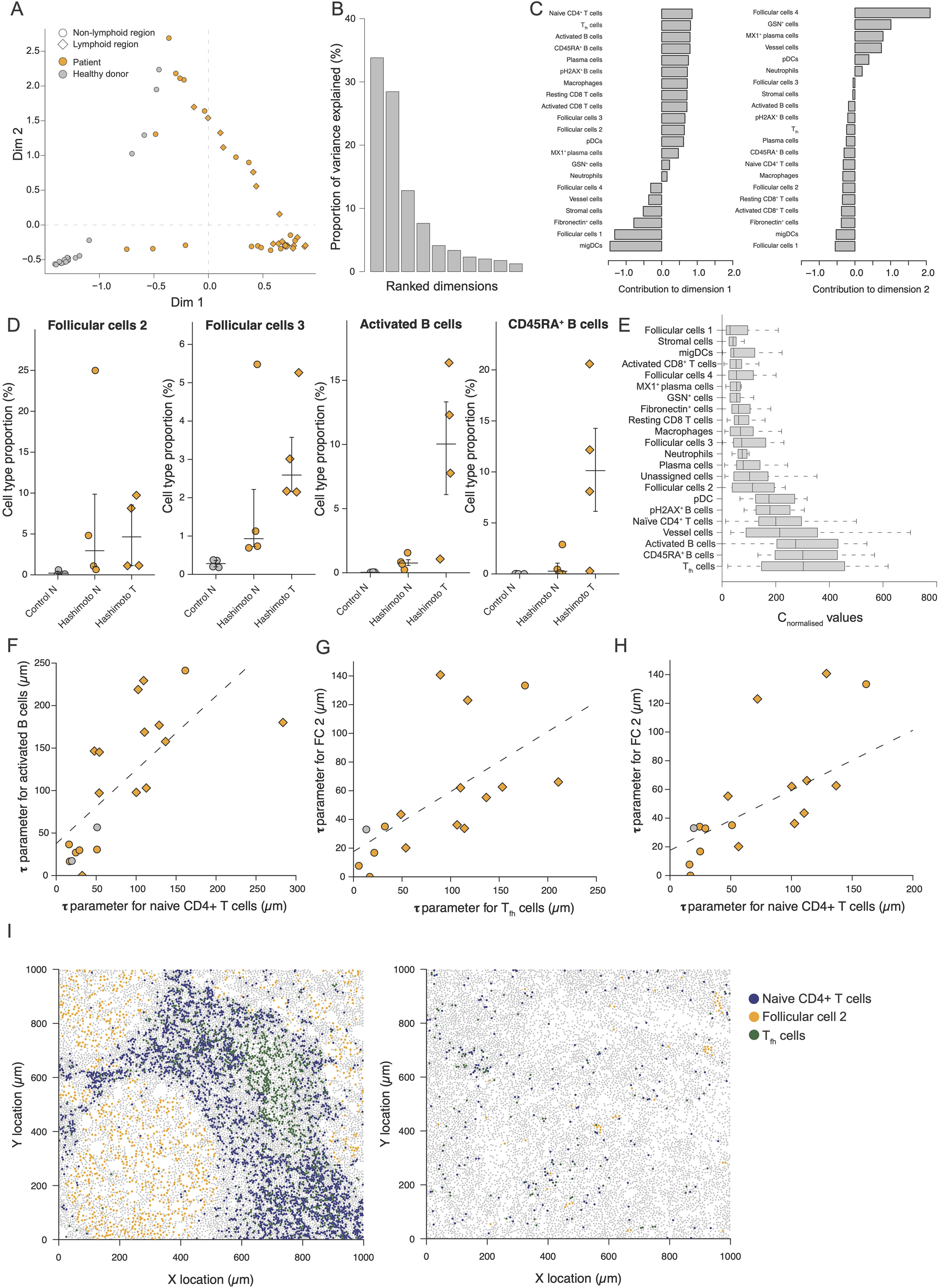
(**A**) Scatter plot of the first two dimensions associated to the Correspondance Analysis of the thyroid dataset. **(B)** Scree plot associated to the Correspondance Analysis of the thyroid dataset. **(C)** Contribution of each cell cluster to the first two dimensions of the Correspondance Analysis. **(D)** Proportion of Follicular cells 2 (left), Follicular cells 3 (center left), Activated B cells (center right) and CD45RA+ B cells (right) across samples. **(E)** Distribution of the C_normalised_ parameter value across cell clusters. **(F)** Comparison of the **τ** parameter for naive CD4+ T cells and Activated B cells. Dashed line corresponds to a regular linear regression. **(G)** Comparison of the **τ** parameter for T_fh_ cells and Follicular Cells 2. Dashed line corresponds to a regular linear regression. **(H)** Comparison of the **τ** parameter for CD4+ T cells and Follicular Cells 2. Dashed line corresponds to a regular linear regression. **(I)** Spatial distribution of cells belonging to the naive CD4+ T cells (blue), Follicular cells 2 (orange) and T_fh_ cells (green) clusters in two representative samples.

**Supplementary Figure 6.**
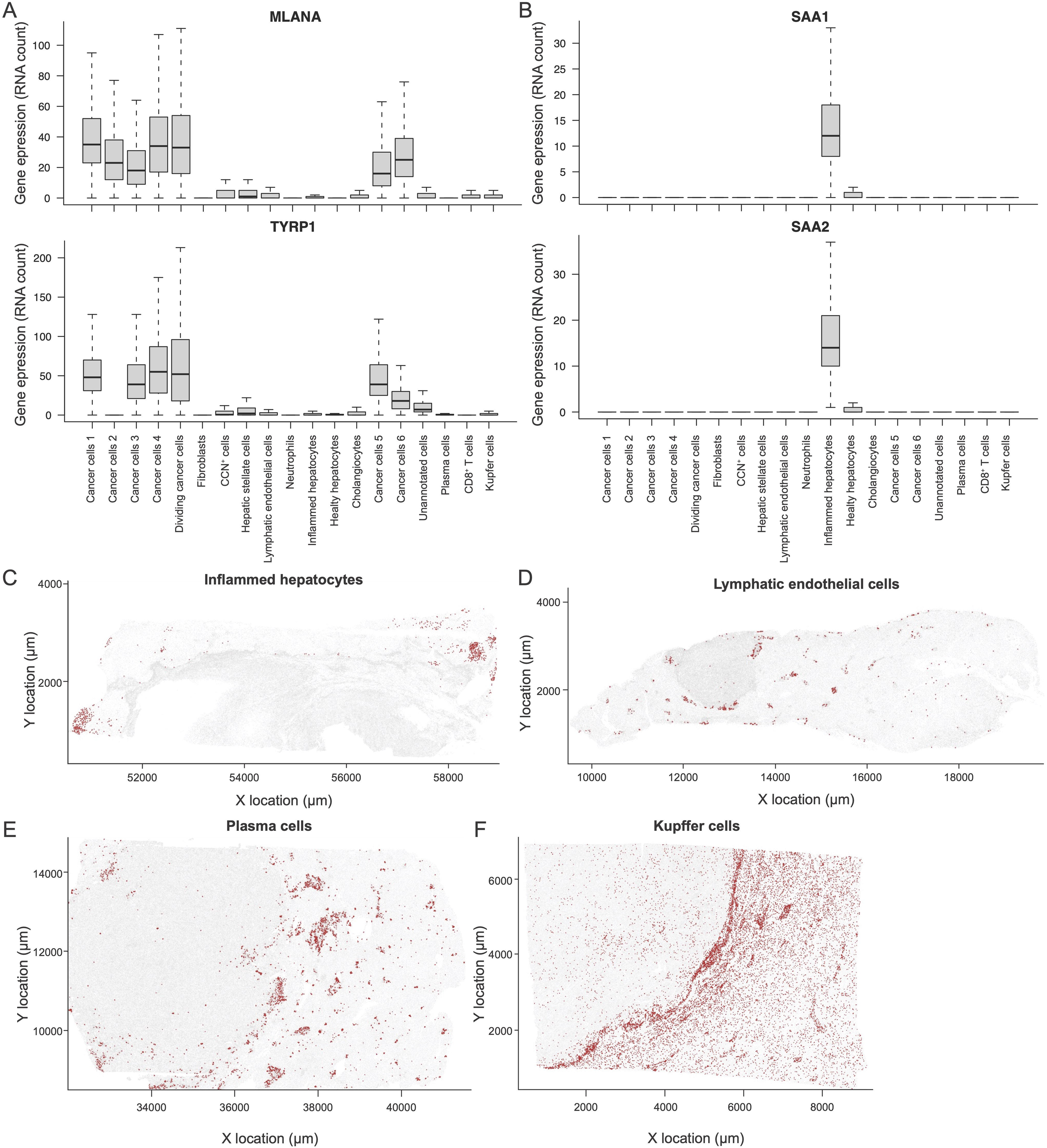
(**A**) Upper panel: Expression of the MLANA gene across cell types. Lower panel: Expression of the TYRP1 gene across cell types. **(B)** Upper panel: Expression of the SAA1 gene across cell types. Lower panel: Expression of the SAA2 gene across cell types. **(C)** Spatial distribution of inflamed hepatocytes in a representative sample. **(D)** Spatial distribution of Lymphatic endothelial cells in a representative sample. **(E)** Spatial distribution of plasma cells in a representative sample. **(F)** Spatial distribution of Kupffer cells in a representative sample.

**Supplementary Figure 7.**
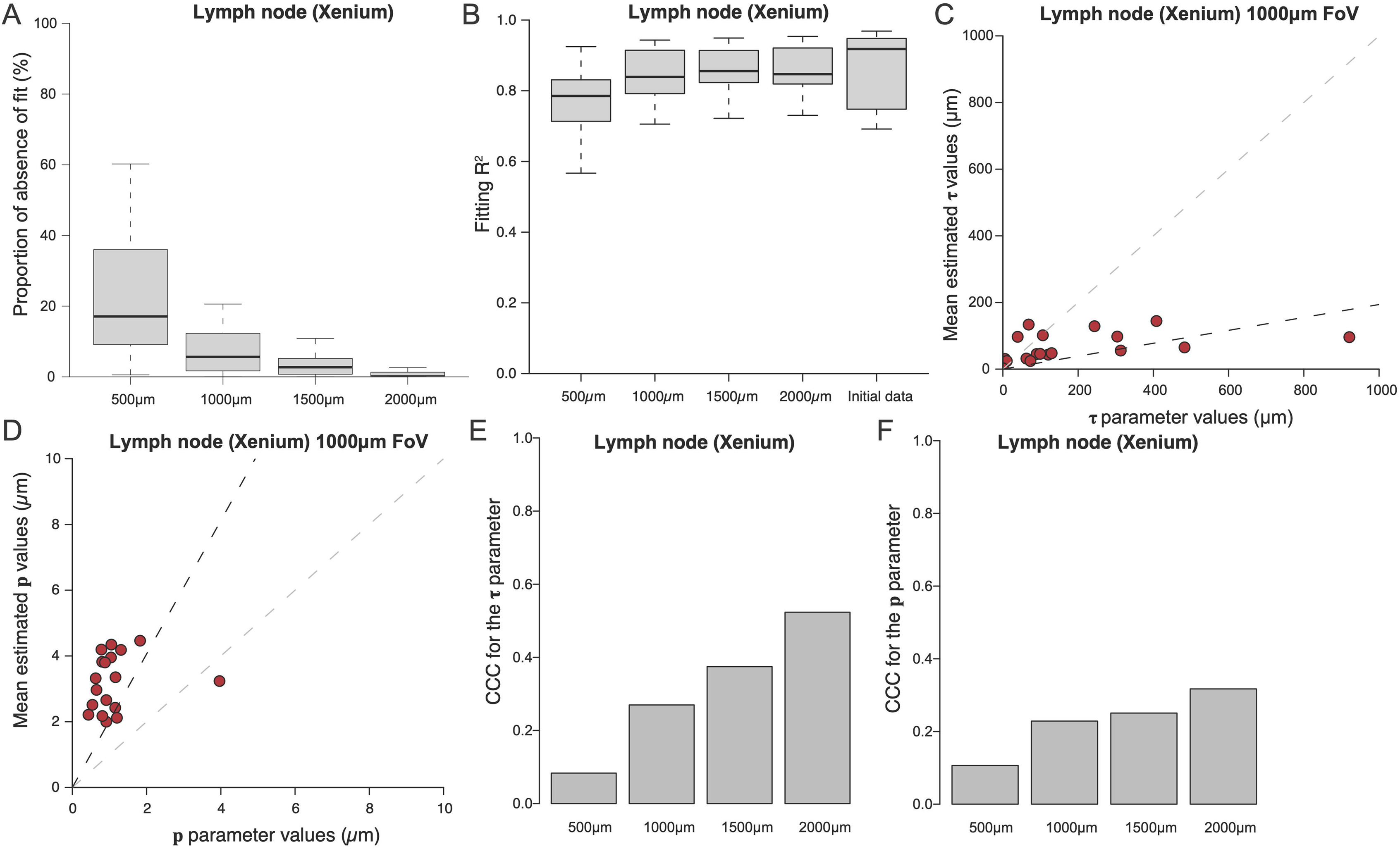
(**A**) Distribution of the proportion of absence of fit across cell types for different FoV size for the Xenium Lymph Node dataset. The boxplot is based on N = 18 cell types. **(B)** Distribution of the R^2^ fit across cell types for different FoV size for the Xenium Lymph Node dataset. The boxplot is based on N = 18 cell types. **(D)** Comparison of the initial **τ** parameter estimation and of the average **τ** estimate using 1000µm FoV on the Xenium Lymph Node dataset. The black dashed line corresponds to the linear fit between the two variables. The grey dashed line corresponds to the line x=y. **(E)** Comparison of the initial **p** parameter estimation and of the average **p** estimate using 1000µm FoV on the Xenium Lymph Node dataset. The black dashed line corresponds to the linear fit between the two variables. The grey dashed line corresponds to the line x=y. **(F)** CCC between between the **τ** parameter values and the average **τ** estimate across the sampled FoVs of different size. **(G)** CCC between between the **p** parameter values and the average **p** estimate across the sampled FoVs of different size.

**Supplementary Table S1:** IMC antibody panel for the analysis of thyroid samples.

**Supplementary Table S2:** Description of UM patient.

**Supplementary Table S3:** Xenium gene panel used for the analysis of UM liver metastasis samples.

## Author contribution

F.M performed the computational analysis and wrote the R code. R.C. designed the thyroid IMC antibody panel and performed the thyroid IMC experiment. K.B. performed UM Xenium experiment. L.D.K. designed the UM cohort and contributed to the design of the project. M.H., C.P and S.D. collected and revised thyroid histological samples. S.R.R designed the UM cohort. B.B. supervised the IMC thyroid experiment. M.R. designed the UM cohort and secured fundings for the Xenium experiment. P.B. developed the methodology, oversaw the project, performed the thyroid IMC experiment, wrote the manuscript and secured fundings.

## Acknowledgments

We acknowledge Bost lab members for critical reading and providing feedback on the manuscript and thank Mengze Zhang, José Francisco Carreño Martinez, Alice Balfourier, Uria Mor and Ronan Thibaut for their valuable advices. F.M. and P.B. were funded by a Junior Principal Investigator Starting grant from Institut Curie.

